# *In vivo* microstimulation with cathodic and anodic asymmetric waveforms modulates spatiotemporal calcium dynamics in cortical neuropil and pyramidal neurons of male mice

**DOI:** 10.1101/2019.12.16.878892

**Authors:** Kevin C. Stieger, James R. Eles, Kip A. Ludwig, Takashi D.Y. Kozai

## Abstract

Electrical stimulation has been critical in the development of an understanding of brain function and disease. Despite its widespread use and obvious clinical potential, the mechanisms governing stimulation in the cortex remain largely unexplored in the context of pulse parameters. Modeling studies have suggested that modulation of stimulation pulse waveform may be able to control the probability of neuronal activation to selectively stimulate either cell bodies or passing fibers depending on the leading polarity. Thus, asymmetric waveforms with equal charge per phase (*i.e.* increasing the leading phase duration and proportionately decreasing the amplitude) may be able to activate a more spatially localized or distributed population of neurons if the leading phase is cathodic or anodic, respectively. Here, we use two-photon and mesoscale calcium imaging of GCaMP6s expressed in excitatory pyramidal neurons of male mice to investigate the role of pulse polarity and waveform asymmetry on the spatiotemporal properties of direct neuronal activation with 10 Hz electrical stimulation. We demonstrate that increasing cathodic asymmetry effectively reduces neuronal activation and results in a more spatially localized subpopulation of activated neurons without sacrificing the density of activated neurons around the electrode. Conversely, increasing anodic asymmetry increases the spatial spread of activation and highly resembles spatiotemporal calcium activity induced by conventional symmetric cathodic stimulation. These results suggest that stimulation polarity and asymmetry can be used to modulate the spatiotemporal dynamics of neuronal activity thus increasing the effective parameter space of electrical stimulation to restore sensation and study circuit dynamics.

**Significance Statement:** Electrical stimulation has promise to restore sensation and motor function, as well as treat the symptoms of several neurological disorders. However, the mechanisms responsible for the beneficial effects of stimulation are not fully understood. This work supports modeling predictions by demonstrating that modulation of the stimulation waveform dramatically affects the spatial recruitment and activity level of neurons *in vivo*. These findings suggest that stimulation waveform symmetry represents a parameter that may be able to increase the dynamic range of stimulation applications. Further characterization of these parameters with frequency, and amplitude could provide further insight into the mechanisms of electrical stimulation.

## 1. Introduction

Implantable stimulating and recording microelectrodes have been fundamental to our understanding of the nervous system. The ability to perturb the brain with electrical stimulation has allowed us to realize localized functions of different brain areas[1] in addition to the functions of specific neural circuits[2, 3]. Additionally, intracortical microstimulation (ICMS) can provide sensory feedback in neural prosthetics and potentially restore sight to the blind [4–6]. Although the use of electrical stimulation has provided critical knowledge to our understanding of the brain, and can relieve symptoms of some neurological disorders [7–9], our poor understanding of the variability in neuronal recruitment and behavioral responses to different stimulation parameters has prevented us from fully understanding the brain and how circuits change in disease[10].

Neuronal representation of sensory information is highly complex [11], and the ability to selectively recruit neuronal populations proximal to the electrode by varying pulse amplitude, frequency, or polarity, could provide the ability to stimulate neural activity that matches physiological response patterns [12–24]. For example, neuronal adaptation is a common response to sustained stimuli and is believed to allow the brain to better respond to transient stimuli [25]. Therefore, the ability to selectively modulate neuronal adaptation to electrical stimulation could help to produce more naturalistic and complex neuronal activity [14]. Furthermore, current stimulation parameters have poor selectivity, and neuronal recruitment depends on the complex order of cells dendrites and axons in the surrounding environment [26, 27].

Electrical stimulation often activates axon nodes or terminals as opposed to cell bodies due to the higher excitability, thus the local axonal projections greatly influence the spatial distribution of activated neurons [16, 28–30]. Additionally, neuronal entrainment to stimulation depends on the site of activation on the neuron, the afterpotential dynamics, and the relative amount of excitatory and inhibitory input to the cells [23, 31, 32]. Our group recently published a study demonstrating that frequency can also modulate the spatial pattern of neuronal activation [24]. Specifically, distal neurons fail to entrain to frequencies ≥ 30 Hz, resulting in a dense population of neurons activated near the electrode. On the other hand, 10 Hz stimulation resulted in steady and spatially distributed neuronal activation. Therefore, consideration of the role of the neuronal element being activated (site of activation; *e.g.* soma, axon, dendrite), its afterpotential dynamics, and baseline excitability, in addition to the neuronal geometry around the electrode during stimulation, could help inform our ability to encode complex sensory information with ICMS.

Modeling [33–35] and *in vivo* [27–29, 36] studies have demonstrated that orthodromic activation of cells and antidromic activation of passing fibers have similar thresholds when using monophasic stimulation pulses [21]. Therefore, it has been proposed that modulation of the likelihood of a neuronal element being activated by electric field may change the neuronal response to electrical stimulation [21, 22, 32]. Although monophasic waveforms have lower activation thresholds, convention is to use charge balanced biphasic waveforms to prevent electrode degradation; however, there can be safety benefits to slight charge imbalance [37]. Within these safety constraints, biphasic electrical stimulation pulse waveforms can be designed by altering the polarity of the leading or activation phase and/or the shape of each phase while maintaining an equivalent charge injection (so that the net injected charge is zero, or slightly reduced in the balance phase). These waveforms can, therefore, take advantage of the nonlinear temporal and voltage dynamics of sodium channel activation to impact the ability to excite different neuronal elements.

McIntyre and Grill [21, 22] investigated the effect of waveform asymmetry on the compartmental excitation of neurons. Specifically, they modeled the effect of increasing the duration and proportionately decreasing the amplitude of the leading phase of charge balanced waveforms on preferential activation of cell bodies or passing axons. Their studies concluded that charge balanced asymmetric waveforms could selectively activate either cell bodies (cathodic first), or passing fibers (anodic first), depending on the leading polarity. Furthermore, proximal, or distal neurons had higher entrainment probabilities depending on leading polarity and frequency [22]. These predictions were supported with an *in vivo* study by Wang and colleagues [38], however they only investigated the effect of a single pulse, while clinical ICMS typically uses trains of pulses. By modulating the activity of the neurons using different stimulation waveform shapes during trains of stimulation, both neuronal entrainment and spatial recruitment could be selectively regulated, thus providing potential for more control over spatiotemporal dynamics of neuronal recruitment.

The present study aims to further characterize the role of stimulation waveform in neuronal activation to better understand the mechanisms underlying electrical stimulation using mesoscale, and two-photon calcium imaging. This study sought to answer two main questions, namely near the safe maximum charge injection 1) Can stimulation waveform modulate the spatial distribution of neuronal activation? And 2) Do different populations of neurons prefer or entrain better to different stimulation waveforms? We address these questions in 4 sections using 30s stimulation trains delivered at 10 Hz to layer II/III of mice densely expressing GCaMP6s in Thy-1 positive neurons [39]. Using mesoscale and two-photon imaging, we demonstrated that the relative activation magnitude, and spatial distribution of neuronal activation can be modulated by the waveform asymmetry, depending on leading polarity. Specifically, increasing the asymmetry of stimulation reduced or increased the area of activated neuronal elements (Section 3.1-3.2) without significantly impacting the density of neurons activated around the electrode (Section 3.3), for cathodic or anodic leading polarity, respectively. Furthermore, at the same current density, asymmetric cathodic pulses were significantly less effective at activating neurons, while asymmetric anodic pulses were comparable to symmetric cathodic stimulation (Section 3.4). The findings of this study ultimately support the claim that stimulation waveform can be an important tool in the modulation of neuronal activation, and highlights the importance of considering the nonlinear dynamics of ion channels when designing electrical stimulation protocols for better control of neuronal activation.

## 2. Methods

### 2.1 Animals and electrode implantation

Seven male, transgenic mice C57BL/6J-Tg(Thy1 GCaMP6s)GP4.3Dkim/J (>8 weeks, >28g, Jackson Laboratories, Bar Harbor, ME) were used in this study[39]. Subjects (n = 5-6 biological replicates) were housed in 12h light/dark cycles with free access to food and water. Acute surgery and electrode implantation followed a protocol previously published [24, 40–49]. Mice were initially anesthetized with ketamine/xylazine (75mg/kg;7mg/kg) intraperitoneally (IP) and updated with ketamine (40 mg/kg) as needed (~ every hour). Mice were then placed on a heating pad and in a stereotaxic frame with continuous O_2_ supply. After removing the scalp, a stainless-steel bone screw (ground and reference electrode) was placed over the motor cortex to minimize the interaction of the electric fields at the working and reference electrodes for a monopolar setup. A dental drill was used to perform bilateral craniotomies (9-12mm^2^) over the somatosensory cortices. Acute style 16-channel single shank Michigan style functional silicon probes with electrode site surface area of 703 μm^2^ (A1×16-3mm-100-703; NeuroNexus, Ann Arbor, MI) were used for all stimulation experiments. Prior to implantation, iridium electrode sites were activated to increase charge storage capacity, and 1kHz impedance was verified to be <500 kOhm [50]. Electrodes were implanted using an oil-hydraulic Microdrive (MO-81, Naishige, Japan) at a 30° angle into the somatosensory cortex to a final depth of 200-300 μm (layer II/III because there is high expression of GCaMP6s in this model [39]). To aid vascular visualization in two-photon experiments, sulforhodamine (SR101; 0.1 mg) was injected IP. Saline was used to keep the brain hydrated throughout the experiments.

### 2.2 Stimulation paradigms

Stimulation was performed using a TDT IZ2 stimulator controlled by an RZ5D system (Tucker-Davis Technologies, Alachua, FL). Asymmetric pulses were designed such that any change in pulse width had a proportionate decrease in amplitude to maintain charge balance to the precision of the stimulator. An asymmetry index was defined similarly to Wang *et al.* 2012 [38], as the ratio of the leading phase duration to the return phase duration minus 1 (1). For example, a symmetric biphasic pulse would have an asymmetry index of 0. All durations and amplitudes of the stimulation pulses are shown in Table 1. The control stimulation is symmetric cathodic-first with 200 μs pulse width and 12.5 μA resulting in 2.5 nC/phase charge injection and 0.36 mC cm^−2^ charge density. This charge density is within the Shannon limits, (k = [−0.06 −0.03]), however, because the site diameter is well below 200 μm (~15μm), the 4 nC/phase safety limit is more appropriate [51, 52]. Thus, these stimulation parameters are within suggested safety limits while being near the limit of maximum safe charge injection [51–55]. Stimulation trials included a 30s baseline followed by a 30s stimulation duration and, either a 20s, or 60s post stimulation period for two-photon, or mesoscale imaging respectively [24]. Stimulation parameters will be referred to by the letter of the leading polarity with a subscript of the asymmetry index (*e.g.* symmetric cathodic = C_0_; See Table 1). For each stimulation session, paradigms were applied in a random order.

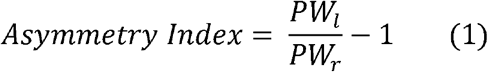

**Table 1:** Stimulation parameters.

2.5nC/phase, 0.36 mC/cm<sup>2</sup>
| Asymmetry Index | Leading Amplitude, μA | Leading Pulse Width, μs | Return Amplitude, μA | Return Pulse Width, μs | Total Pulse Time, μs |
| --- | --- | --- | --- | --- | --- |
| -1/2 | $\pm 12.5$ | 200 | $\mp 6.2$ | 400 | 1800 |
| 0 | $\pm 12.5$ | 200 | $\mp 12.5$ | 200 | 1200 |
| 4 | $\pm 2.5$ | 1000 | $\mp 12.5$ | 200 | 400 |
| 7 | $\pm 1.6$ | 1600 | $\mp 12.5$ | 200 | 600 |

### 2.3 Mesoscale and two-photon imaging

Mesoscale imaging was performed during stimulation over a 3.7×3.0 mm window using an MVX-10 epifluorescence microscope (Olympus, Tokyo, Japan) and a CCD camera with a 50 ms exposure time (Retiga R1, QI imaging) at 20 Hz. For cellular resolution, a two-photon microscope (Bruker, Madison, WI) with an OPO laser (Insight DS+, Spectra Physics, Menlo Park, CA) tuned to 920 nm and kept below 40 mW was used for neuronal and neuropil imaging at 30 Hz using resonance mode. The microscope was equipped with a 16X 0.8 NA water immersion objective (Nikon Instruments, Melville, NY) with a 3mm working distance resulting FOV of 407 × 407 μm^2^ (512 × 512 pixels). Image acquisition was initiated by a trigger from the TDT system.

### 2.4 Image analysis and statistics

#### 2.4.1 Stimulation inclusion

All analyses contained 5 or 6 biological replicates for mesoscale or two-photon imaging, respectively. Each stimulation session contained one train of each of the 8 waveforms delivered at 10 Hz. Separate sessions were carried out at different stimulation sites and considered independent as shown by Histed and colleagues [16]. Cells and vasculature were used to verify minimal tissue displacement during each stimulation trial so that the same cells could be identified throughout each session. Trials were excluded if there was observable tissue displacement.

#### 2.4.2 MVX GCaMP6s Quantification

All analysis of the MVX images was performed in MATLAB (MathWorks, Boston, MA). The images were initially spatially smoothed using a gaussian filter with 2 standard deviations, then a 2×2 mm^2^ ROI was selected around the stimulation probe. Fluorescence was transformed into ΔF/F_0_ using the mean fluorescence over the 30s pre-stimulus baseline. Each pixel was then filtered using a low-pass finite-impulse response filter (FIR) with a cutoff frequency of 2 Hz [56]. To quantify the spatial spread of GCaMP activity, ΔF/F_0_ was calculated in 20 μm radial bins and transformed to activation area (Fig. 1). Activation area is defined as the area of the maximal activated bin such that all smaller bins were also activated. A bin was considered activated if its ΔF/F_0_ was greater than the mean plus 2 standard deviations of the baseline for 1s consecutively (Fig. 1). This harsh thresholding aided in removing motion artifacts but did not alter the quantification of activation. The magnitude of activation area was quantified over time to determine how far neural elements can be from the electrode during maximal and sustained activation. One animal was excluded in the calculation of percent change in area because activation area was 0 mm^2^ in the first 5 s.

**Figure 1:**
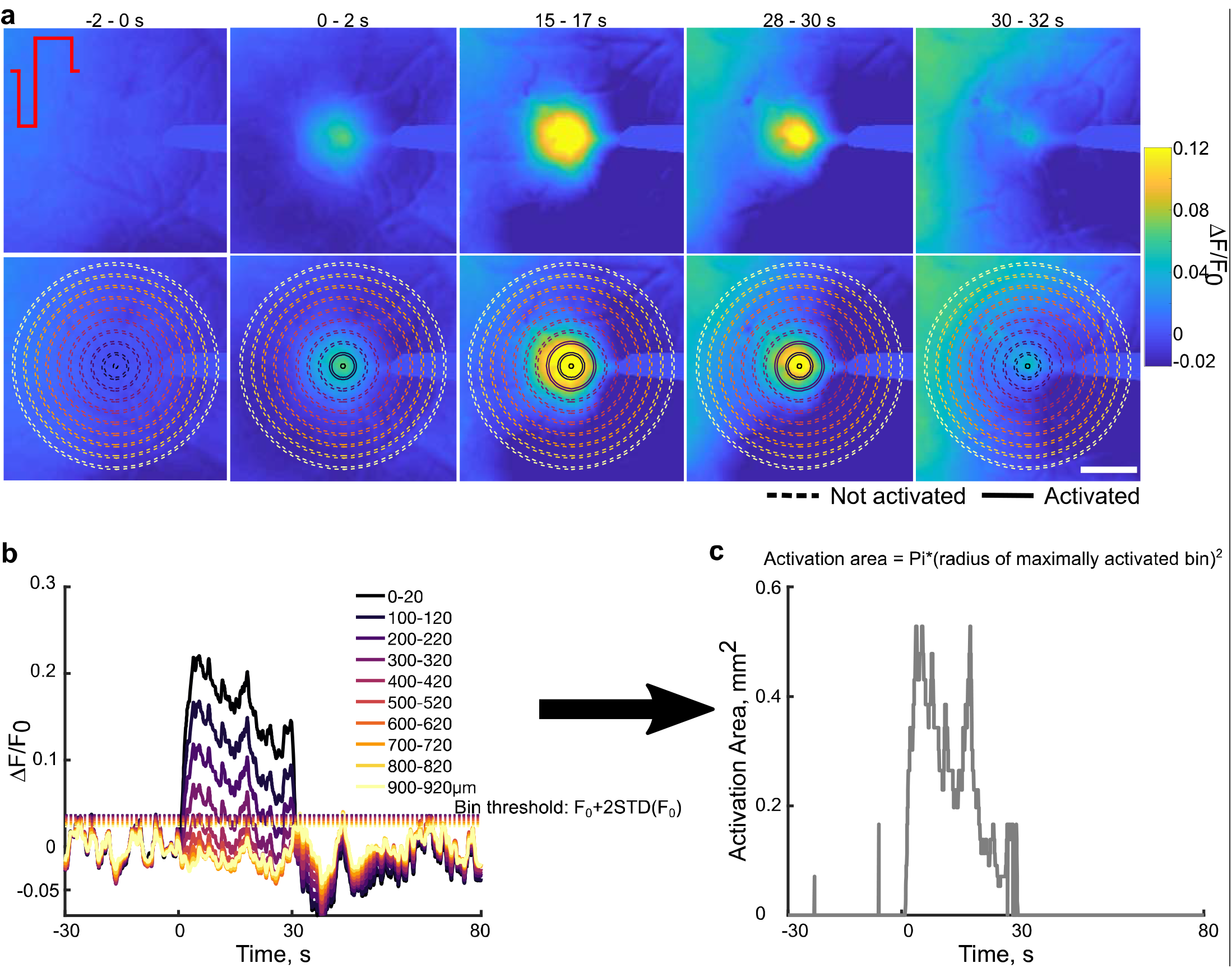
Quantification of activation area. (a) top: Representative ΔF/F_0_ images over the stimulation show that over time the spread of activation and intensity increases then decreases by the end of stimulation. Bottom: The same images in the top overlaid with a sub selection of radial bins used to calculate the fluorescence intensity and activation area. Solid lines represent activated bins. Scale bar is 500 μm. (b) The fluorescence intensity in the radial bins plotted in (a) with each bin’s corresponding activation threshold (dotted lines). (c) Using the radius of the maximally activated bin with all smaller bins also activated, the activation area was calculated over time and shows an increase then a decrease over time.

The magnitude of fluorescence change was determined to quantify the intensity of neural activity by averaging over the first 0.98 mm. This distance threshold (0.98 mm) was defined as the minimum bin where the mean stimulation intensity of each stimulation condition was less than the grand mean plus one standard deviation of the mean (over 30s) at the maximum bin (1.4mm). Then, the mean, peak, and slope of activation were calculated using the average fluorescence of this area for each stimulation condition. To calculate the slope of activation or inactivation, a first order polynomial was fit to the activation over the first or last two seconds of stimulation.

#### 2.4.3 Neuronal soma quantification

For each stimulation session, neurons were manually identified using the 2D projection of the fluorescence standard deviation over the 30s stimulation and outlined using the ROI manager in ImageJ (NIH)[24]. The mean fluorescence intensity of each cell was extracted for further analysis in MATLAB. The fluorescence intensity for each neuron was then temporally filtered as was done with the MVX images. Neurons were considered activated if the ΔF/F_0_ was greater than F_0_ plus three times the standard deviation of the baseline period for at least 1 s. The distance of all activated cells was determined and placed in 20 μm radial bins to determine the spatial distribution and density of neurons activated by each stimulation condition.

To distinguish the timing of activation, cells that maintained calcium activity for the first 2 s but not the last 2 were classified as “onset responsive”, cells that were active during the last 2 s but not the first 2 seconds were considered “delayed responsive” and all other cells were considered “steady state responsive” [24]. The “activation time” was calculated as the time for the fluorescence to reach half the peak cumulative fluorescence over the 30 s, and represents the time it takes to reach half the response energy, as done previously[24]. The peak, mean, time to peak and slope of activation of neurons were determined similarly to the MVX activation.

To quantify how specific neurons respond to different waveforms, we first determined which cells were activated in common between the different parameters. Next, an activation index ranging from − 1 to 1 (Equation 2) was created to quantify how specific neurons respond to different stimulation conditions. Where X_1_, and X_2_ represent the two stimulation conditions being compared. Statistical significance was determined by performing a Student’s t-test to determine if the activation index was significantly different from 0.

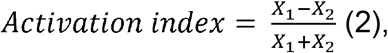

#### 2.4.4 Neuropil Quantification

To quantify changes in fluorescence of the neuropil, the electrode and all identified neuronal cell bodies were masked out (see section 3.4.3) [24, 57, 58]. Similar to section 3.4.2, the changes in neuropil fluorescence as ΔF/F_0_ were quantified in 20 μm radial bins radiating form the electrode site. Quantification of activation area and fluorescence magnitude metrics were calculated as was done in section 3.4.2. One animal was excluded from the time to peak area analysis because there was no area of activation for the C_4_ condition. Fluorescence intensity quantifications were performed over all imaged bins to a distance of 260 μm from the electrode site.

#### 2.4.5 Statistical analysis

All data in violin plots (Bastian Bechtold 2016, Violin plots for Matlab, https://github.com/bastibe/Violinplot-Matlab/blob/master/violinplot.m) show the mean (horizontal line), the median (diamonds), grey box plots, (25^th^ and 75^th^ percentiles, 1.5 times the interquartile range), scatter plot of the data within the kernel density estimate, with outliers as black “+”. In order to determine the interaction of asymmetry and polarity on outcome measures statistical analysis in violin plots was assessed with a two-way repeated measures ANOVA (2-way rmANOVA; alpha = 0.05, including outliers), because the stimulation parameters in a trial were measured on the same neurons. This analysis was performed in R (R Core Team (2019). R: A language and environment for statistical computing. R Foundation for Statistical Computing,Vienna, Austria. URL https://www.R-project.org/.) using the ez package (Michael A. Lawrence (2016). ez: Easy Analysis and Visualization of Factorial Experiments. R package version 4.4-0.https://CRAN.R-project.org/package=ez). Significance was assessed for neuron density and neuron number over distance (Fig. 6ab) using a three-way repeated measures ANOVA (polarity, asymmetry, distance). Lastly, Student’s t-tests were used to determine activation indices significantly deviating from 0 (Fig. 8). Box plots show the mean (diamonds), the median (vertical line) and 25^th^ and 75^th^ percentiles, plus 1.5 times the interquartile range (whiskers). Outliers are represented with a black “+”. In cases where sphericity was violated, the Greenhouse-Geisser method was used to correct degrees of freedom. Post-hoc analysis consisted of pairwise unequal variance (Welch’s) t-tests with a Holm-Bonferroni correction. Data are described as mean ± standard deviation (STD).

## 3. Results

Stimulation waveform asymmetry was predicted to modulate the spatial selectivity of neuronal activation [22]. Here, we used *in vivo* mesoscale and two-photon calcium imaging in Thy1-GCaMP6s mice, which highly express GCaMP in layer II/III, to investigate how the stimulation waveform and polarity modulate the magnitude of neural activation and spatial distribution of activated neurons and neuropil. We delivered 30s stimulation trains at 10 Hz using a single shank Michigan-style electrode implanted into layer II/III of somatosensory cortex. Although higher frequencies are used in clinical ICMS [4], 10 Hz stimulation was employed to study the effect of stimulation waveform on the spatial pattern of activation, while decoupling the spatial fall off of neural activation that occurs at higher frequencies [24]. All stimulation parameters delivered 2.48-2.56 nC/phase charge injection and 353-364 μC cm^−2^ charge density for 30s following a 30s baseline, and are compared to conventional symmetric cathodic first square wave with 200 μs pulse width and 12.5 μA amplitude similar to parameters used in human clinical ICMS applications (See Table 1)[4, 6, 59]. Stimulation waveform significantly influenced the spread of neuronal activation as well as intensity of activation under two-photon and mesoscale imaging, without impacting the density of activated neurons proximal to the electrode. Lastly, we quantified how specific neurons responded to different waveforms and show that the majority of neurons were most strongly activated by cathodic stimulation with long return phase (*i.e.* second), and anodic stimulation with long activation phase (*i.e.* first), but the populations of activated neurons were largely overlapping.

### 3.1 Mesoscale imaging reveals strong interaction of stimulation polarity and asymmetry on the spread of cortical pyramidal neuron activation

To determine the impact of waveform shape on the spatiotemporal spread of activation on a large spatial scale, mesoscale imaging was employed to readout Ca activity during ICMS (Fig. 2). The spread of activation increased radially with limited bias towards any specific direction (Fig. 2a); therefore, the activation area was calculated by determining the area of the most distal activated 20 μm radial bin (ΔF/F_0_> F_0_ + 2 STD) such that all smaller bins were also activated (Fig. 1, Fig. 2b). Interestingly, activation area often peaked within the first 10 seconds and gradually decreased over the duration of stimulation (Fig. 2b). As a measure of the spread of activation at the onset, the peak area was quantified to represent the maximum spread of activation, where greater peak area could suggest greater antidromic activation or indirect activation through monosynaptic connections for short duration stimulation trains (Fig. 2c). As asymmetry index was increased, peak activation area decreased for cathodic-first stimulation and increased for anodic first asymmetric pulses. Repeated measures two-way ANOVA revealed a significant interaction of polarity and asymmetry (n=5, p=0.0017). This is consistent with previous studies suggesting that pulse asymmetry modulates the probability of activating neurons antidromically, where a larger area would suggest greater antidromic activation [21, 38]. Quantification of the time to peak area was comparable across our parameters, suggesting that the spread rate of the initial activation may not be significantly influenced by pulse shape when delivered at 10 Hz (Fig. 2d).

**Figure 2:**
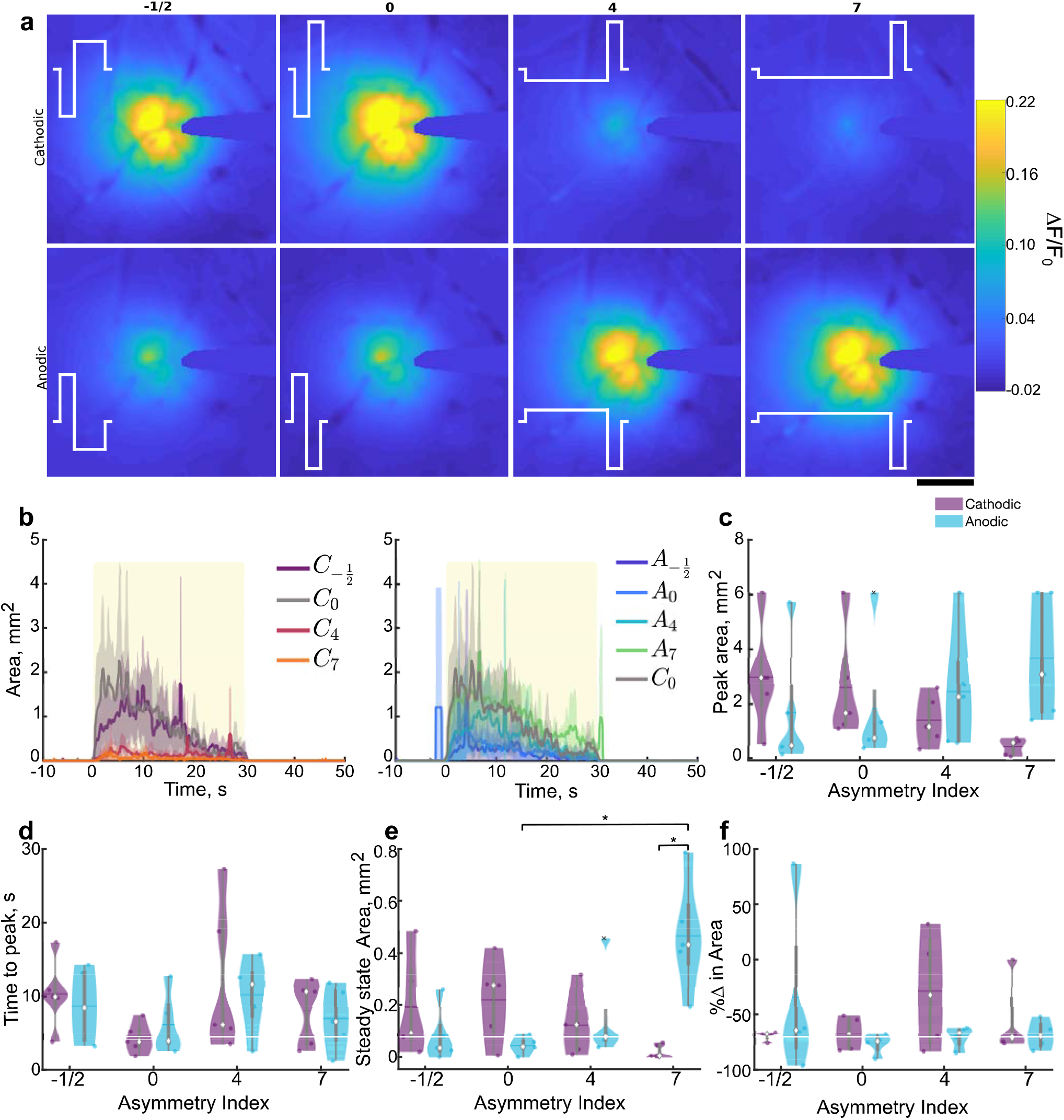
Stimulation polarity and asymmetry affect the spread of cortical activation during 30s stimulation trains under mesoscale imaging. (a) Representative mean ΔF/F_0_ over the 30s stimulation for the 8 different stimulation conditions. The stimulation waveform is represented in white, and the corresponding asymmetry index is displayed above. Scale bar indicates 500 μm. (b) Average (±SE; n=5) cortical activation area peaks early on (within 10-12s) and decreases over time and is comparable between C_−1/2_, C_0_, A_4_, and A_7_ and is reduced in C_4_, C_7_, A_−1/2_, and A_0_. Yellow box represents stimulation. (c-d) Peak (c; n=5, F_3,12_= 9.54, p=0.0017) and time to peak (d) area during the stimulation train, where the median is indicated as a diamond and the mean indicated as a horizontal line in the violin plots. (e) Steady state area is increased in A_7_ compared to A_0_ and C_7_ (n=5, F_3,12_=10.36, p=0.0012). (f) Area rises and reduces over time resulting in a negative percent change in area for all stimulation conditions. Statistics were first assessed with a 2-way rmANOVA followed by post-hoc Welch’s T-tests with Holm correction. * indicates p<0.05.

Next, in order to quantify stable spread of activation during long stimulation trials, steady state area was calculated as the average activation area during the last 5s of stimulation, where the activation area stabilized (Fig. 2b,e). Steady state area followed the same pattern of activation as peak area, with asymmetric anodic-first stimulation having significantly greater area compared to symmetric anodic activation, and cathodic first asymmetric pulses (post-hoc Welch’s T-test p<0.05; Fig. 2e). Despite the expectation that neuronal entrainment to 10 Hz stimulation would be high [24, 60], most stimulation asymmetries resulted in >50% reduction in the activation area between the first and last 5s of the train (Fig. 2f). Together, these data support the hypothesis that stimulation polarity and asymmetry may be able to selectively modulate the spatial spread of activation where cathodic first asymmetric pulses preferentially activate cell bodies and anodic first asymmetric pulses activate passing fibers.

To determine if different populations of neurons prefer, or entrain better to, particular stimulation waveforms, we investigated interaction of stimulation polarity and asymmetry on intensity of activation by quantifying the magnitude of fluorescence changes over time and distance (Fig. 3). Similar to activation area, measuring the ΔF/F_0_ in radial bins revealed a transient activation that peaked within the first few seconds (Fig. 3a, c; S1) and gradually reduced before plateauing by the end of stimulation (Fig. 3a). However, the ΔF/F_0_ appeared more stable for cathodic first asymmetric pulses with a long leading phase (Fig. 3a). Furthermore, averaging the ΔF/F_0_ over the 30s stimulation revealed an exponential decay with distance that became comparable by 980 μm from the electrode (Fig. 3b; see methods). Therefore, the ΔF/F_0_ was averaged over the first 980 μm from the electrode to compare across stimulation conditions. This averaging did not change the observed pattern of activation (Fig. 3c).

**Figure 3:**
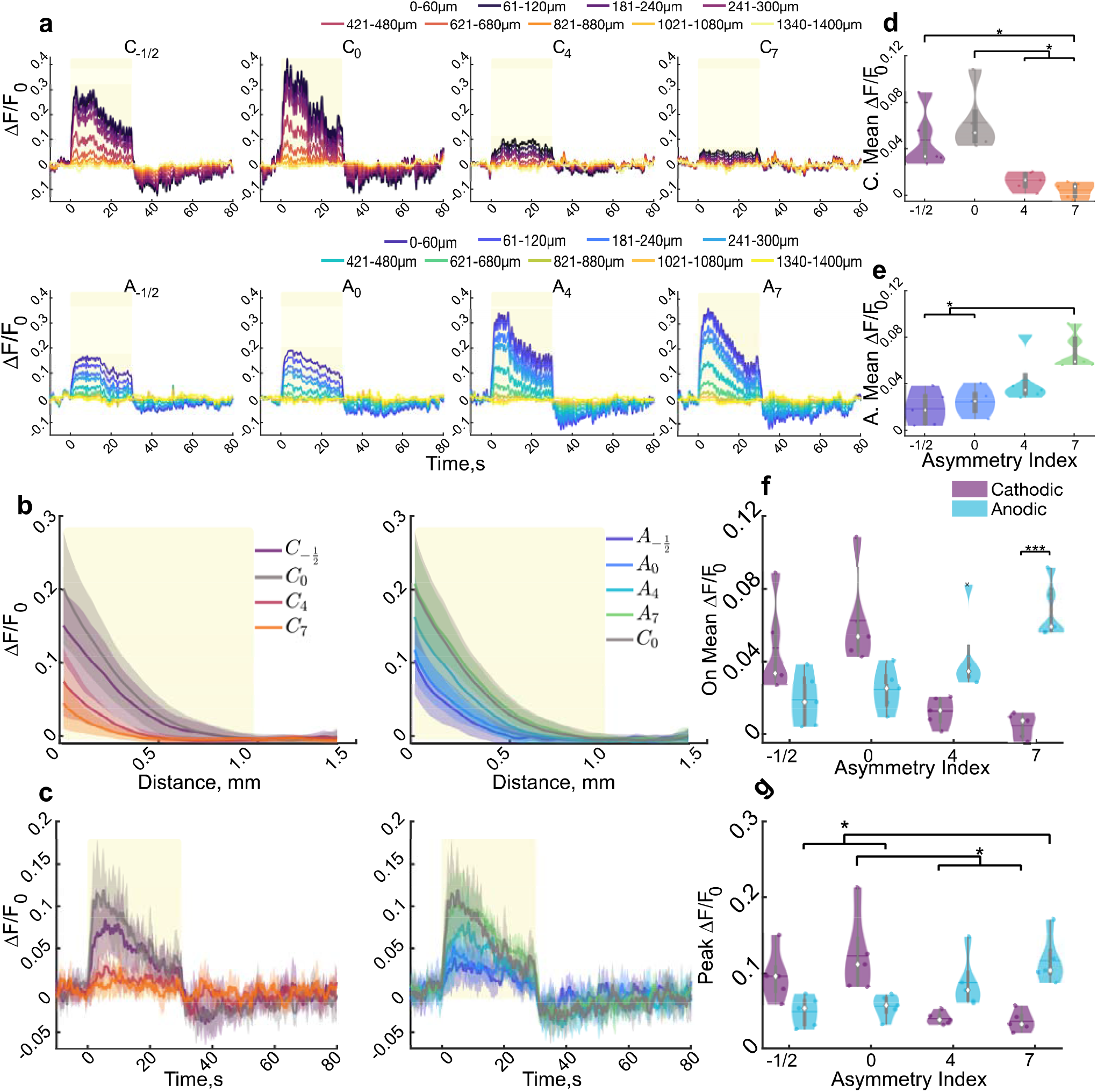
Stimulation pulse waveform and polarity modulate cortical neuronal activity under mesoscale imaging. (a). Cortical activity peaks within the first 15 seconds and reduces over time and distance for a representative animal then is inhibited for several seconds following stimulation. (b) the average (±STD) neuronal activity over the stimulation train reduces exponentially with distance from the electrode and is reduced in C_4_ and C_7_. Yellow rectangle indicates the distance (980μm) which will be averaged over for all but (k). (c) The average ± STD activation over 980μm follows the same pattern as the binned intensities. (d-f) Mean activation intensity over the stimulation period for the cathodic (d), anodic (e) and the interaction (f; F_3,12_=30.87, n=5, p<1e−5). Cathodic and anodic were separated for clarity to show all post-hoc significance. (g) Peak activation intensity decreases or increases as cathodic or anodic asymmetry increases, respectively (F_3,12_=14.36, p<0.001). Statistics were first assessed with a 2-way rmANOVA followed by post-hoc Welch’s T-tests with Bonferroni-Holm correction. : * indicates p<0.05, ** indicates p<0.01, ** indicates p<0.001.

Next, the peak and mean fluorescence were determined to quantify the maximal and sustainable magnitude of neuronal activation during stimulation trials, respectively (Fig 2d-g). This quantification revealed a significant interaction between asymmetry and polarity for mean (p<1e+5) and peak (p<0.001) ΔF/F_0_. Increasing cathodic asymmetry resulted in a significantly reduced mean and peak ΔF/F_0_ compared to symmetric cathodic stimulation (Fig. 3 d-f). On the other hand, anodic-first pulses with the longest duration leading phase (A_7_) resulted in an increased mean and peak activation intensity compared to symmetric anodic stimulation (Fig. 3 d-f).

Furthermore, quantification of the rate of change of fluorescence during the stimulation onset or termination revealed a significant interaction of polarity and asymmetry for the activation (Fig. S1; F_3,12_=9.16, n=5, p=0.002). Lastly, several stimulation conditions were followed by a substantial and sustained reduction below baseline, localized within ~400 μm around the electrode (Fig. 3a, c). Averaging over the 10s following the stimulation revealed a significant interaction of stimulation asymmetry and polarity on the sustained reduction in ΔF/F_0_ (Fig. S1). These results are consistent with the idea that, at the same current density, neuronal activation threshold is comparable for anodic first asymmetric pulses, and is increased for cathodic first asymmetric pulses [21]. Taken together, these results suggest that stimulation polarity and asymmetry can selectively modulate spatial localization of neuronal activation around the electrode, and the magnitude of activation is similarly affected.

### 3.2 Two-photon imaging demonstrates strong interaction of polarity and asymmetry on spread of neuropil activation

In order to distinguish calcium activity between neuron somas and neuropil, ICMS was repeated under two-photon microscopy. First neuropil calcium dynamics were quantified to better describe the effective spread of activation (Fig. 5)[61]. Figure 5a shows the standard deviation images of a stimulation session without neurons masked out to highlight how the neuropil activation relates to neuronal activation. While mesoscale imaging revealed a peak within the first 10 s (Fig. 3b), followed by a gradual decline, neuropil activation (within 260μm from the electrode site) was often more stable during stimulation, consistent with our previous report [24] (Fig. 4a-b). Therefore, the mean activation area was quantified in addition to the peak and steady state (Fig. 4c-e). There was a significant interaction of polarity and asymmetry (F_3,15_=18.16, n=6, p<1e−4) on the steady state area, where increasing asymmetry for cathodic pulses reduced activation area, and increasing asymmetry for anodic pulses increased the area (Fig 3c). This effect was also present for mean (Fig. 4d; F_3,15_ = 76.3, n=6, p<1e−8) and peak area (Fig. 4e; F_3,15_=34.91, p<1e−6). Furthermore, the time to peak area was elevated for cathodic stimulation with long activation phases (C_4_, C_7_) and anodic stimulation with symmetric phases or an increased return phase (A_0_, A_−1/2_) (Fig. 4f). Post-hoc analysis revealed an increased time to peak activation area for cathodic pulses with increased activation phase duration (C_4_, C_7_) compared to the other cathodic parameters (C_0_, C_−1/2_) suggesting that the activation of neurons could be increasing during the stimulation train.

**Figure 4:**
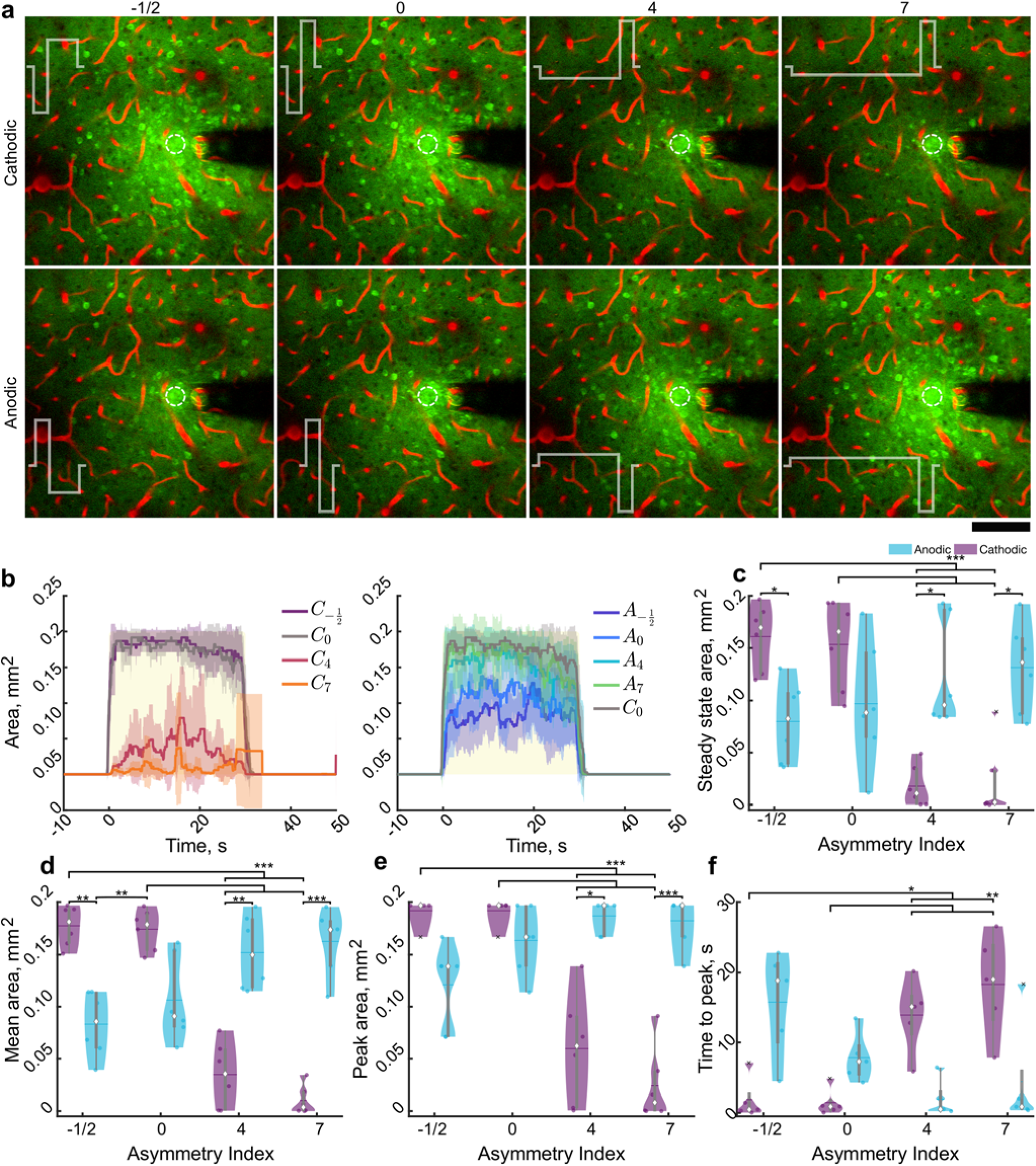
Stimulation polarity and asymmetry modulate the activation area of neuropil under two-photon imaging. (a) Representative standard deviation images over the stimulation train suggest stimulation waveform and asymmetry spread and intensity of neuropil (and neuronal activation). Stimulation waveform is represented in white, and the asymmetry index is displayed above the images. Scale bar represents 100 μm. (b) Average (±STD, n=6) neuropil activation area peaks early on after stimulation and is largely stable during stimulation and is reduced in C_4_, C_7_, A_−1/2_ and A_0_ compared to C_0_, yellow box represents stimulation. (c-d) Stimulation polarity, asymmetry and the interaction significantly modulate the steady state (c; F_3,15_=18.16, n=6, p<1e−4) average (d; F_315_=76.31,n=6, p<1e−8), peak (e, F_3,15_=34.9, n=6, p<1e−6), and time to peak (f, F_3,12_=15.90, n=6, p<0.001) activation area over time. Statistics were first assessed with a 2-way rmANOVA followed by post-hoc Welch’s T-tests with Holm correction. * indicates p<0.05, ** indicates p<0.01, *** indicates p<0.001.

**Figure 5:**
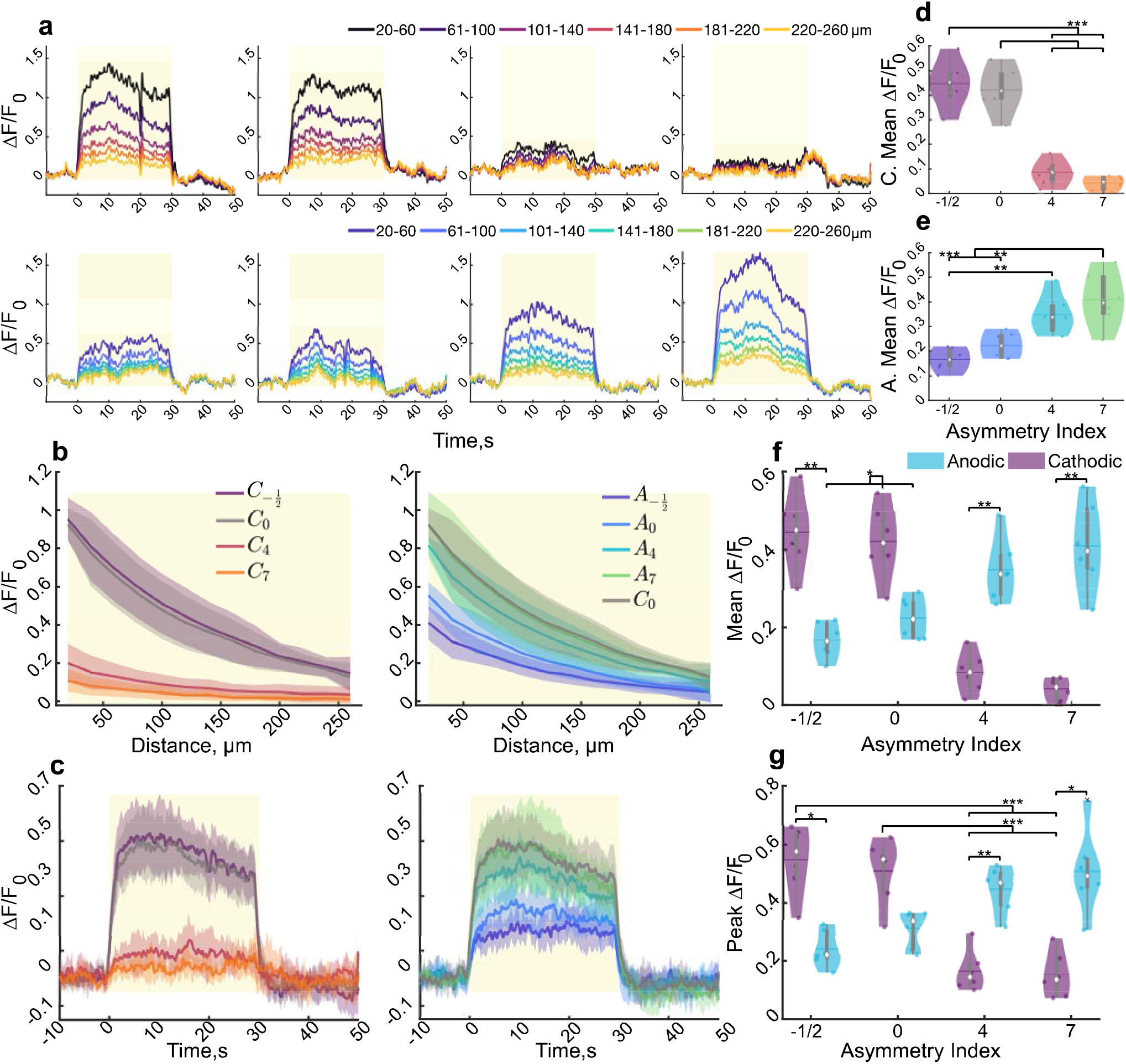
Stimulation polarity and waveform modulate the level of neuropil activation over time. (a) Representative ΔF/F_0_ in increasing radial bins during stimulation demonstrate differential effects of neuropil activation intensity with polarity and asymmetry. Yellow box represents stimulation. (b) The average stimulation over time (± STD) decreases exponentially over distance and appears reduced in C_4_, C_7_, A_−1/2_, A_0_ compared to C_0_. Yellow box represents area to be averaged over for comparisons. (c) The average neuropil activation intensity (±STD) over 260μm rises rapidly then slowly increases followed by a slow decrease over time. (d-f) The average stimulation intensity of the neuropil in the imaging plane over the stimulation train is significantly reduced in C_4_ and C_7_ Compared to C_−1/2_ and C_0_, and is reduced in A_−1/2_, and A_0_ compared to A_7_ (F_0.97,4.88_=48.60, n=6, p<1e−4). (g-j) Peak (g, F_0.93,4.64_=45.06, n=6, p<0.001), time to peak (h, F_3,15_=5.06, n=6, p=0.0128), activation (I; F_3,15_=24.47, n=6, p<1e−5) and inactivation slope (j; F_0.70,3.52_=21.15, n=6, p=0.0049) during stimulation averages over 260 μm. Statistics were first assessed with a 2-way rmANOVA followed by post-hoc Welch’s T-tests with Holm correction : ** indicates p<0.01, *** indicates p<0.001.

Next, the spatiotemporal dynamics of the calcium transients in the neuropil were examined over the imaging plane in order to evaluate activation preference to different stimulation waveforms (Fig. 5). The activation intensity remained relatively constant during stimulation while decaying exponentially with distance, although at a lesser degree than was observed under mesoscale imaging (Fig. 5a-b, 2b). Consistent with mesoscale imaging, cathodic stimulation with either symmetric phases or increased return phase (C_0_,C_−1/2_) showed comparable neuropil entrainment with anodic first asymmetric stimulation with a longer activation phase (A_4_, A_7_), which were elevated compared to the other stimulation parameters (Fig. 5c). Quantification of the mean (Fig. 5d-f), peak (Fig. 5g), and time to peak activation (Fig. S2), as well as the rate of fluorescence change over the first and last two seconds of stimulation (activation/inactivation slope; Fig. S2) revealed trends consistent with mesoscale imaging. Specifically, neuropil entrainment was reduced as asymmetry increased for cathodic stimulation and entrainment increased for anodic stimulation as quantified with the mean (interaction: F_0.96,4.88_=48.6, n=6, p<1e−4), peak (interaction: F_0.93,4.64_=45.06, n=6, p<0.001), activation (F_3,15_=24.47, n=6, p<1e−5) and inactivation slope (F_0.7,3.52_=21.25, n=6, p<0.0049), in addition to the time to peak fluorescence (F_3,15_=5.06,n=6, p=0.013). Together, neuropil activation dynamics agree with the mesoscale analysis and suggest that the stimulation waveform likely plays a role in both the spatial spread of neuronal activation, as well as the magnitude of activation.

### 3.3 Two-photon imaging of neuronal activation reveals consistent interaction of polarity and asymmetry on spatial recruitment and neuronal activity

We next aimed to determine the spatial preference of neural activation to different waveforms by characterizing the neuronal somatic activation over distance (Fig. 6). After determining which neurons had fluorescence intensities 3 standard deviations above the mean for at least one second over the 30s train, the neuron’s distance from the electrode was used to calculate the activation density. Neuronal density was calculated as the number of neurons in the area of circles with increasing radii. All stimulation conditions resulted in dense neural activation that was significantly greater within the first 40μm compared to the area beyond 80 μm (Fig. 6a). Furthermore, the mean distance of activated neurons was significantly modulated by stimulation polarity and asymmetry (Fig. 6c; interaction: F_3,15_=27.84, n=6, p<1e−5). Specifically, the mean distance of activated neurons reduced as cathodic asymmetry increased and generally increased as anodic asymmetry increased. This interaction was also observed for the total number of neurons activated (F_0.78,3.89_=17.60, n=6, p=0.0053; Fig. 6c). These data are consistent with Figures 2-4 and further support the hypothesis that asymmetric cathodic leading stimulation may selectively activate neurons orthodromically, and anodic first stimulation may be more selective for antidromic activation of passing fibers.

**Figure 6:**
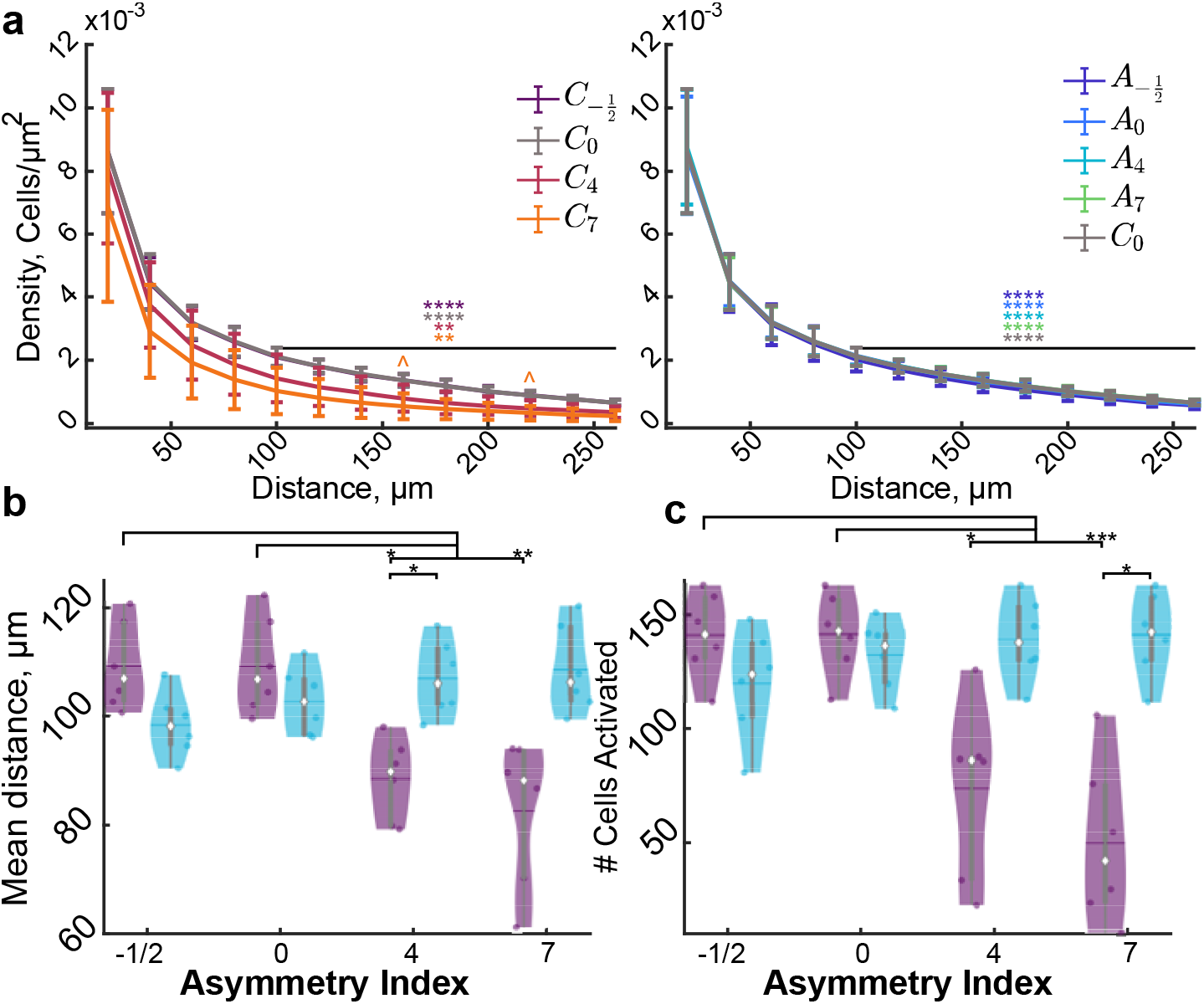
Stimulation asymmetry and polarity modulate the spatial distribution of neuronal activation. (a) The density of activated neurons within the first 40 μm of the electrode is significantly greater than the density of neurons beyond 40 μm in all conditions except C_7_ (F_5,36_ = 3.77, n=6, p<1e−10). (b) The number of activated cells is activated in relatively even amounts within all conditions except C_4_ and C_7_ where the majority of activated neurons ~70% is within the first 106μm. Data represented as mean ± STD (F_5,3_=4.02, n=6, p<0.012). (c-d) Stimulation polarity and waveform modulate the mean distance of the activated neurons (F_3,15_=27.84, n=6, p<1e−5) as well as the number (F_0.78,3.89_=17.60, n=6, p<0.0053). Statistics were first assessed with a 3-way rmANOVA (a,b) or a 2-way rmANOVA (c,d) followed by post-hoc Welch’s T-tests with Bonferroni-Holm correction * indicates p<0.05, ** indicates p<0.01, *** p<0.0001, **** p<1e−5, ^ indicate comparison to C_0_ p<0.05.

Next, the intensity level of neuronal activation was quantified to determine how the relative neuronal activation differed for these stimulation waveforms (Fig. 7). There was a dynamic range of neuronal responses to stimulation (Fig. 7a-b). In general, quantification of neuronal activation was consistent with the mesoscale imaging and neuropil analysis, but with larger effect size. There was a highly significant interaction of polarity and asymmetry on the mean, activation and inactivation slope (Fig. S3), peak, and time to peak fluorescence (all p<0.001). Specifically, neurons were not activated as strongly to cathodic first asymmetric stimulation with increased activation phase duration, but gradually increased fluorescence (Fig. 7c-g). On the other hand, neurons activated by anodic first asymmetric stimulation with a long activation phase had highly comparable activation patterns with conventional symmetric cathodic stimulation. These results are consistent with the modeling by McIntyre and Grill, suggesting that the activation threshold for cathodic asymmetric stimulation would be higher than symmetric, and anodic-first stimulation would be comparable [21].

**Figure 7:**
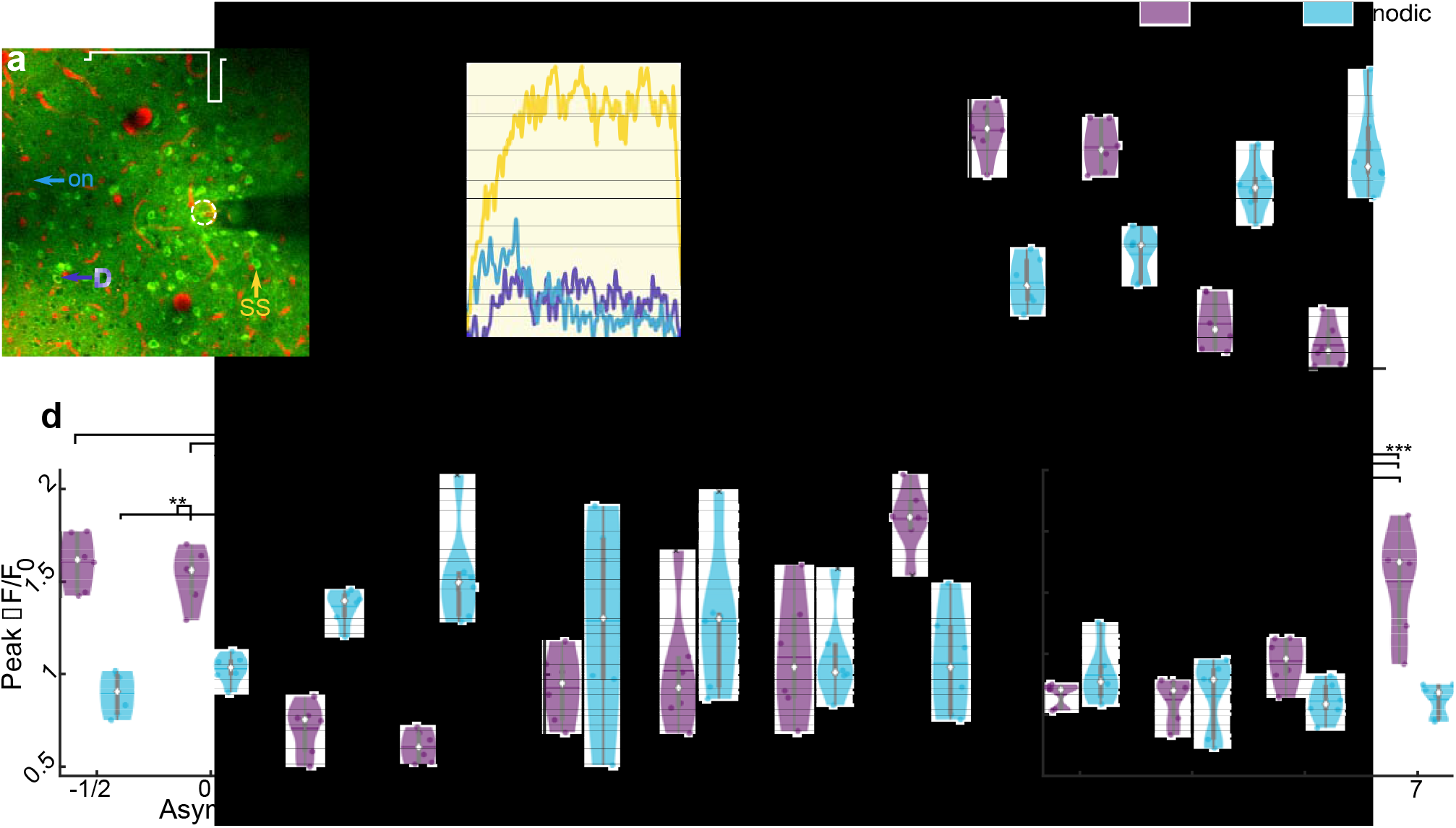
Stimulation asymmetry and polarity modulate neuronal activity. (a) Representative standard deviation image of neurons activated by A_7_. (b) Representative traces of cells indicated in (a) for onset (On), steady state (SS), and delayed (D). Yellow rectangle indicates stimulation. (c-g) Mean (c, F_3,15_=61.86, n=6, p<1e−7), peak (d; F_0.95,4.74_=56.40, n=6, p<1e−4) and time to peak activation (e; F_3,15_=11.10, n=6, p<0.001) of cells averaged for each animal. (k) Average weighted activation time of activated cells for each animal (F_3,15_=10.96, n=6, p<0.001). Statistics were first assessed with a 2-way rmANOVA followed by post-hoc Welch’s T-tests with Holm correction : * indicates p<0.05, ** indicates p<0.01, ** indicates p<0.001.

Finally, the activation time was quantified as defined by Michelson et al, as the time for the cumulative sum of ΔF/F_0_ to reach 50 % of the total sum[24]. In this definition, a neuron whose activity peaks and is maintained from 0-30s, will have an activation time of 15 s. There was a significant interaction of stimulation polarity and asymmetry on the activation time, where increasing cathodic asymmetry resulted in a greater activation time (Fig. 7k). Consistent with this, although the percentage of steady state cells were the majority for all waveforms, both the percentage of steady state cells (active for the first and last two seconds) and delayed response cells (active for the last two seconds, but not the first two) were significantly modulated by stimulation asymmetry and polarity (Fig. S3) Overall, these data suggest stimulation waveform asymmetry can modulate the spatial selectivity of neural activation, with the potential cost of reducing activation magnitude, on average.

### 3.4 Stimulation polarity and asymmetry modulate population of activated neurons

Although, on average, neurons were most strongly activated by symmetric cathodic, asymmetric cathodic with a long return phase and anodic with a long activation phase, there are some cells uniquely activated by the different stimulation conditions (Fig. 8a; white arrows, red cells); however, the populations were largely overlapping (yellow cells and neuropil). Some neurons appeared to prefer certain stimulation waveforms as shown by greater calcium transients (Fig. 8 a-c). Therefore, the overlap in the neuronal populations activated by the different waveforms and their activation properties were measured in order to quantify the selectivity and preference different populations of cortical neurons around the electrode. While, Calculating the percent overlap in the populations of neurons activated by different waveforms revealed that while asymmetric cathodic first pulses with a long activation phase can activate a smaller subset of the neuronal population around the electrode (Fig. 8d, Fig. 6), asymmetric anodic stimulation pulses with a long leading phase result in activation of both the neurons in close proximity to the electrode and distal neurons whose fibers of passage pass by the electrode (>80% overlap for all anodic waveforms).

**Figure 8:**
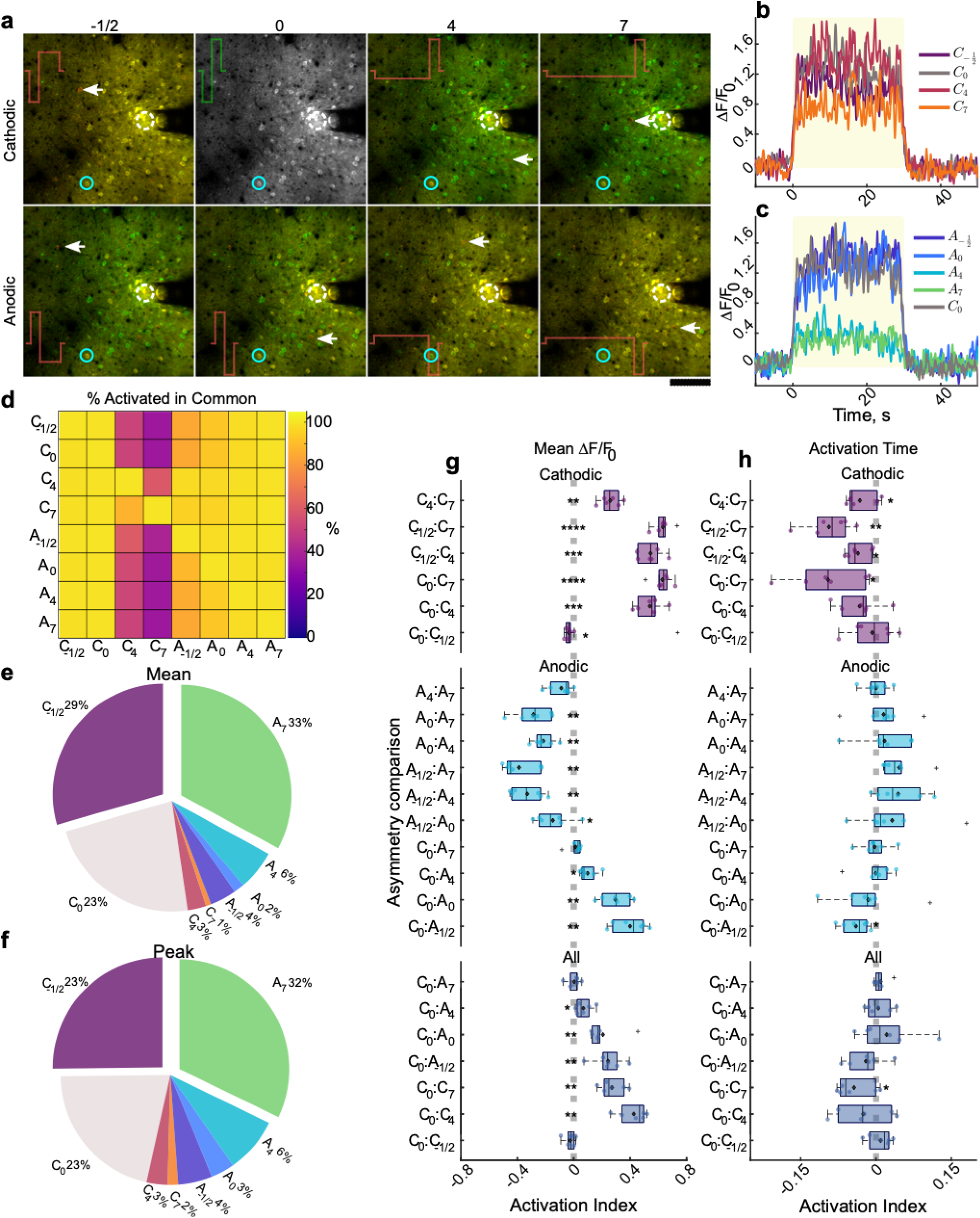
Population of activated cells is largely overlapping between stimulation conditions, and the level of activation is comparable between C_0_, C_−1/2_, and A_7_. (a) Representative composite standard deviation images of the cells activated by conventional C_0_ in green, or each respective stimulation waveform in red. Yellow cells represent neurons activated by both stimulation conditions. Note the relative level of overlap between C_−1/2_, C_0_, A_4_, and A_7_. White arrows signify cells activated by comparison waveform that was not activated by symmetric cathodic stimulation. (b-c) Representative traces of neuron indicated by blue circle in (a) activated by each stimulation waveform. (d) Heatmap of the cells activated in common between the conditions on the vertical and horizontal axis. The values represent the percent of cells activated by the vertical condition that were also activated by the condition on the horizontal axis. Note the nearly 100 % overlap of activation for all conditions except C_4_ and C_7_. (e-f) Pie charts of the relative percent of neurons in the population activated by all conditions that preferred (greatest mean (e) or peak(f) each stimulation condition. (g-h) Activation index of the relative mean (g) or activation time (h) for each comparison where the activation index ranges from −1 to 1 and a positive index indicates greater magnitude of the first condition in the comparisons. The cells quantified in the top, middle, and bottom groups were all activated by all cathodic waveforms (top), all anodic waveforms and symmetric cathodic (middle), or all waveforms (bottom) N=6 for each t-test. * p<0.05, ** p<0.01, *** p<1e−5, ****p<1e−8.

Given that the different neuronal populations around the electrode (proximal vs distal) were predicted to entrain better to the different stimulation waveforms (cathodic asymmetric vs anodic asymmetric) [22], we next aimed to characterize the preference of activation to the different waveforms. In general, of the neuronal population that was activated by all stimulation waveforms, the neurons were activated most strongly by anodic asymmetric stimulation with the longest leading phase (A_7_) and cathodic stimulation with a longer duration return phase (C_−1/2_); however, a small percentage preferred asymmetric cathodic stimulation (Fig. 8 e-f). Quantification of activation preference (See Section 2.4.3) revealed consistent trends with previous figures, where, on average for each animal, the activation index was shifted from center (significant difference in preference) for cathodic stimulation that was symmetric (C_0_) or had longer duration return phase (C_−1/2_), and asymmetric anodic stimulation (A_4_, A_7_). Interestingly, the activation time between conditions was generally more comparable in all groups except among cathodic stimulation parameters (Fig. 8h). Altogether, these data demonstrate that stimulation waveform can substantially impact the activation threshold, for some neurons, but not all.

## 4. Discussion

Given that performance of stimulation can be greatly altered by stimulation parameters, it is critical to identify how stimulation parameters affect the tissue surrounding the electrode. In particular, the spatial distribution, relative spike rate, and number of activated neurons [26, 62-66]. Here, we investigated the role of stimulation polarity and asymmetry delivered at 10 Hz in a monopolar configuration on these properties at two different spatial scales. Specifically, we consider how varying the duration of the activation phase relative to the return phase of the stimulation (positive vs negative asymmetry index, respectively) pulses for both cathodic-first and anodic-first stimulation affects the spatiotemporal activation of neurons around the electrode. Although 10 Hz stimulation is lower than what is typically used in the clinic [4], it allowed us to decouple the spatial effects of stimulation waveform from the high frequency mediated inactivation of distal neurons (within 400 μm)[24]. Both mesoscale and two-photon imaging were employed to better quantify the spatial scale of neuronal activation with different stimulation waveforms. The results are summarized in Table 2, and demonstrate that modulation of the stimulation waveform strongly affects both the spatial localization of neuronal activation, and the relative entrainment of neurons to a 30s stimulation train delivered at 10 Hz. Together these results demonstrate the value of stimulation waveform as a tool to modulate the spatial localization of neuronal activation as well as the effective entrainment of neurons to the stimulus.

**Table 2:**
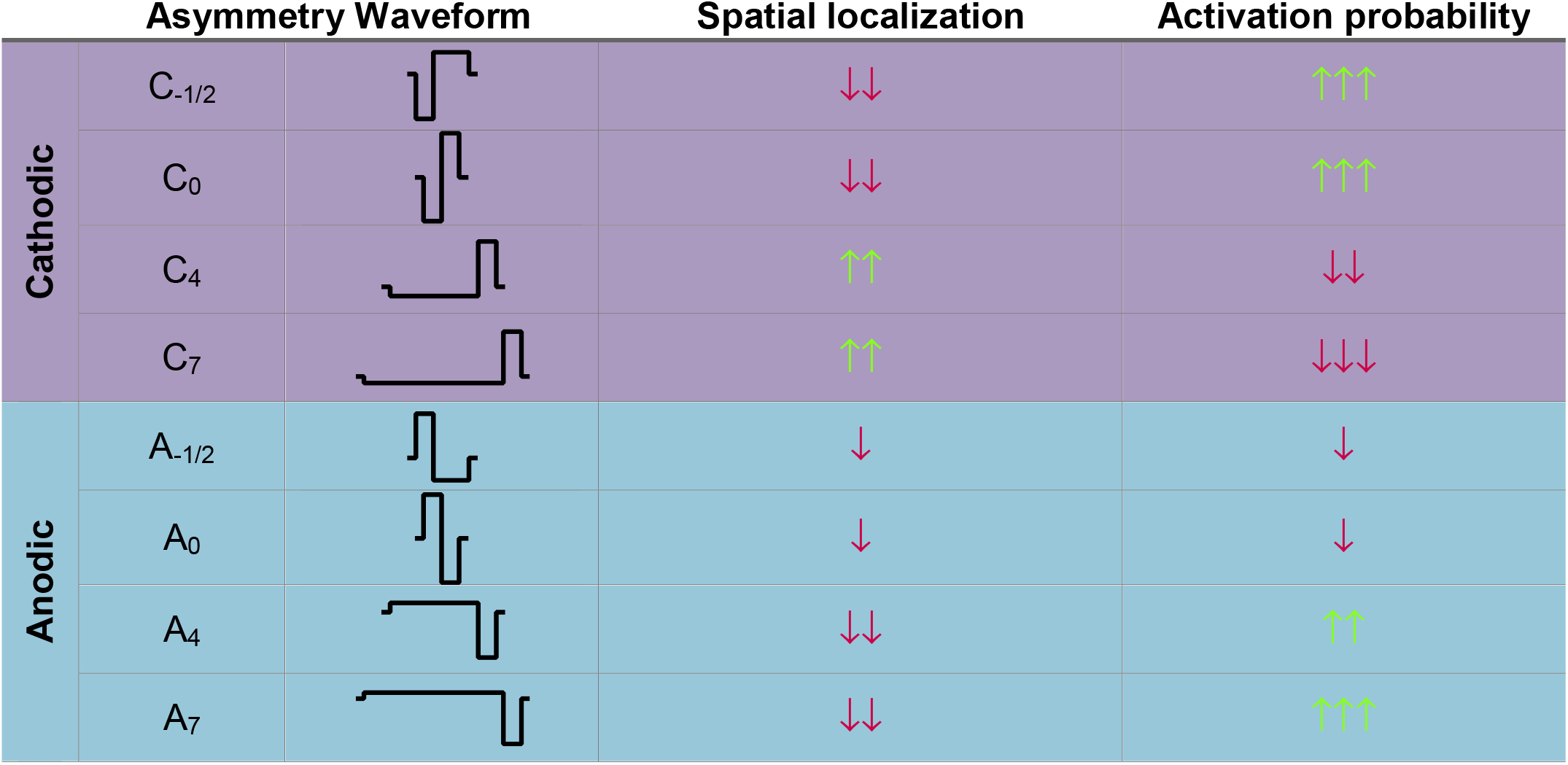
Waveform effects on spatial selectivity and neuronal activation. Waveform effects on spatial selectivity and neuronal activation. Increasing cathodic asymmetry increases spatial localization and reduces neuronal activation. Increasing anodic asymmetry reduces spatial localization and increases neuronal activation. Arrows represent the level of spatial localization or neuronal activation where ↑↑↑ represents the greatest amount and ↓↓↓ represents the least amount.

### 4.1 Biophysics of stimulation pulse waveform on neural activation

Electrical stimulation aims to activate neurons by redistributing ions in the complex tissue geometry around the electrode such that the extracellular environment close to neuronal voltage sensitive sodium channels can become sufficiently negative to depolarize the membrane potential and increase the probability of firing an action potential [67]. In fact, neural activation thresholds are impacted by many pulse parameters including stimulation pulse width, polarity, and pulse shape [21, 27, 28, 68]. Monophasic waveforms can increase the selectivity of neuronal element activation (*i.e.* soma, axon) at lower thresholds than biphasic waveforms [27, 69]. Additionally, shorter pulse durations require exponentially higher amplitudes for effective neuronal activation [28]. This property is often used to characterize the excitability of a neuron to a stimulation pulse [28, 30], where the minimum current amplitude required to elicit an action potential with a long pulse width is called the rheobase current, and the pulse duration required to elicit an action potential at twice the rheobase current is the chronaxie time [28, 35]. However, biphasic waveforms are used to limit electrochemical degradation of the electrode material and subsequent tissue damage [70]. Consequently, there has been an effort to modify the waveform of biphasic pulses to achieve a similar level of activation selectivity as monophasic pulses [21, 71, 72].

#### 4.1.1 Biophysics of cathodic leading stimulation asymmetry on neural activation

It was predicted that increasing the duration and proportionately decreasing the amplitude of the activation phase of stimulation pulses could achieve similar selective activation of different neuronal elements at the cost of increasing the activation threshold [21]. Because the activation function depends on the second spatial derivative of the electric field [35], this may occur by slightly depolarizing off target elements and hyperpolarizing target elements. Specifically, for cathodic leading stimulation, the long duration subthreshold depolarization of off target elements (proximal fibers) aims to increase the number of inactivated sodium channels, while the hyperpolarization of target elements (cell bodies) aims to increase the open probability of the inactivation gate, therefore priming them for activation. For asymmetric cathodic stimulation with a long leading phase, the waveform therefore approximates monophasic anodic stimulation, which is thought to preferentially activate cell bodies, but at higher activation thresholds [21]. Therefore, it would be expected that as the pulse waveform appears more monophasic, the neuronal activation threshold would be affected.

This was confirmed as increasing the asymmetry (*i.e.* pulse width) of cathodic leading pulses reduced the number of neurons activated (Fig. 6), and the activated neurons and neuropil had significantly reduced calcium activity (Fig. 3, 5, 7). Because the current density was kept constant, this would suggest that, on average, the activation threshold is increased for asymmetric cathodic stimulation. Furthermore, increasing the amplitude of asymmetric cathodic stimulation of rat thalamic barreloids achieved a similar magnitude of cortical activation measured by voltage sensitive dye imaging [38]. On the other hand, the magnitude of calcium activity was often strongest for cathodic stimulation with longer return phase (C_−1/2_; Fig. 5, 7, 8). Increasing the duration of the return phase of cathodic stimulation (negative asymmetry index) effectively allows the electric field to persist in the tissue longer. These waveforms would, therefore, approximate monophasic cathodic stimulation and reduce activation threshold [70]. Future studies should investigate the level of neuronal activity as both amplitude and waveform are modulated. Consideration of virtual anodes and cathodes could influence those results; however, there is not much characterization of these effects within the range of safe stimulation amplitudes.

#### 4.1.2 Biophysics of anodic leading asymmetric stimulation on neural activity

Increasing the asymmetry of anodic leading pulses increased the magnitude of neuronal activation to levels similar to symmetric cathodic pulses (Fig. 3,5,7). The mechanism could be similar to anodic break excitation, where the leading anodic phase acts to hyperpolarize the environment putting the sodium channels in an “inactive-activatable” state resulting in activation at the release of the anodic phase [73]. The cathodic phase then acts to increase this effect, thus increasing the depolarizing potential after the termination of the anodic phase. This would be expected to result in reduced activation threshold and increased neuronal activation. Another possibility is that as the anodic asymmetry increases, the stimulation approximates monophasic cathodic stimulation, which is expected to have lower activation thresholds. This could explain the increased activation compared to symmetric anodic leading stimulation (Fig. 3, 5, 7) and strong preference for the A_7_ waveform (Fig. 8). However, it is more likely the leading phase has an impact.

Increasing the duration of the return phase of anodic leading stimulation (A_−1/2_) often resulted in reduced activation magnitudes (Fig. 3, 5, 7). This would further support the influence of the anodic break + cathodic pulse excitation mechanism, since the lower amplitude cathodic return phase would have less depolarizing influence compared to symmetric anodic stimulation, resulting in less neural activity. It is likely a combination of these mechanisms that are modulating the activation thresholds of different neural elements, which could explain the increased behavioral detection thresholds of these waveforms in the rat auditory cortex [18]. However, in cochlear implants, anodic stimulation with long return phase had a lower detection threshold than symmetric anodic stimulation [20]. These differential results further emphasize the complexity of electrical stimulation and highlight the importance of the fiber geometry and organization of excitable elements around the electrode.

#### 4.1.3 Biophysics of the interaction of polarity and asymmetry on neural activity

Cathodic leading stimulation pulses are often used in ICMS because they have lower behavioral detection thresholds compared to anodic leading stimulation [74]. Symmetric cathodic leading stimulation in the guinea pig auditory cortex was shown to elicit greater neuronal responses, particularly in infragranular cortical layers, leading the authors to suggest that cathodic leading stimulation is more effective at activating neurons [68]. Consistent with this, we demonstrate that symmetric cathodic leading stimulation results in greater calcium responses in neuropil and neurons compared to symmetric anodic stimulation even in layer II/III (Fig. 5, 7, 8). Because the activation function depends on the second spatial derivative of the potential [35], cathodic stimulation is likely to most effectively activate neurons and fibers that pass within a close proximity of the electrode (~20 μm [75]), while anodic stimulation may rely on a combination of the formation of a virtual cathode at greater distances from the electrode, or anodic break excitation with a cathodic pulse. However, it should be noted that these results may not be true for all areas of the brain or even considering different neuronal subtypes in the same brain area.

In a modeling study, Anderson *et al*. [12] demonstrated that fibers oriented orthogonally to the electrode had lower activation thresholds to anodic leading stimulation, while passing fibers had lower activation thresholds to cathodic leading stimulation. Furthermore, a separate modeling study investigating surface stimulation polarity demonstrated that, for many cell types, anodic stimulation would have lower activation thresholds [76]. Given the complex fiber orientations in layer II/III, it would make sense that some neurons would respond similarly to cathodic and anodic stimulation. This may explain why some neurons (~2-3%) responded more strongly to symmetric anodic stimulation than any other stimulation condition (Fig. 8). Consistent with this, Voigt *et al.* reported cathodic and anodic stimulation could result in comparable activation in the supragranular layers [68]. This may also explain why 4-5 % of neurons preferred asymmetric cathodic stimulation (Fig. 8) even though the activation threshold was expected to be greater. In fact, it is possible that inhibitory neurons were displaying the unexpected preference [77]. Future studies should investigate the stimulation thresholds (*i.e.* pulse width, amplitude) of different neuronal subtypes given their diverse morphology.

#### 4.1.4 Safety of asymmetric waveforms

Although stimulation waveforms may be charge balanced, it is possible that they are not electrochemically balanced, which can be harmful to both the electrode and the tissue [70]. A long duration activation phase may allow more electrochemical species to dissipate too far from the electrode preventing the charge from being recovered during the return phase, thus resulting in irreversible Faradaic reactions. This may be especially present in trains of pulses, and can be more probable for anodic first pulses. In fact, even increasing the duration of the return phase can reduce the tissue safety [70]. We have observed gas bubble formation under two-photon imaging when stimulating beyond the normal safety limits (indicative of the potential exceeding the water window) [40]; no gas bubble formation was observed in the present study; however, other irreversible reactions could have occurred [70]. Because stimulation waveform has the potential to improve spatial selectivity of neuronal activation it will be important to evaluate the safety of these waveforms for both electrode and tissue damage.

### 4.2.1 Cathodic asymmetry reduces spatial spread of neural activation

The effect of stimulation asymmetry on activation of different neuronal elements has been investigated before in both neuromeric and psychometric analyses [18, 38, 78]. Specifically, using voltage sensitive dye imaging, Wang and colleagues showed that microstimulation of a rat thalamic barreloid with a single pulse of cathodic leading asymmetric stimulation with a long leading phase resulted in a smaller and targeted area of cortical activation, compared to symmetric cathodic stimulation [38]. Additionally, when varying stimulation magnitude, the increase in cortical activation area was smaller in asymmetric cathodic stimulation compared to symmetric cathodic for the same increase in activation magnitude. Therefore, they concluded that asymmetric cathodic stimulation improved the spatial specificity of activation.

Consistent with these results, under mesoscale imaging and two-photon imaging of calcium activity we show that the activation area is significantly reduced for cathodic leading asymmetric waveforms (Fig. 2,4). Furthermore, the mean distance of activated neurons was significantly reduced without significantly impacting the density of neurons activated (Fig. 6). This further supports the hypothesis that asymmetric waveforms with longer activation phases can modulate the activation of fibers or cell bodies. Although, it is possible that the stimulation amplitude of the asymmetric cathodic waveforms could be subthreshold for many neurons [21]. In fact, increasing the amplitude of stimulation can increase the density of neuronal activation at greater distances [75], and smaller amplitudes resulted in less cortical spread as measured with LFPs [79]. Therefore, stimulating with a suprathreshold amplitude, which is likely required for artificial sensation [18, 78], could result in a loss of the spatial selectivity [79]. However, because the cortical activation magnitude increased at a greater rate than the activation area as shown by Wang *et al.,*[38] these waveforms could still maintain spatial selectivity even at higher amplitudes. Future work would need to evaluate the impact of stimulation amplitude and waveform shape, considering both anodic and cathodic leading polarity.

### 4.2.2 Anodic asymmetry increases the spatial spread of neural activation

Anodic leading asymmetric waveforms with a long leading phase were predicted to selectively activate passing fibers, with limited entrainment of proximal cell bodies [22]. Consistent with this, asymmetric anodic stimulation of a thalamic barreloid resulted in greater spatial spread of cortical activation compared to symmetric cathodic stimulation leading the authors to conclude that there was greater activation of passing fibers [38]. In the present study, increasing the asymmetry of anodic leading pulses, significantly increased the spatial spread of neuronal activation under mesoscale (Fig. 2) and two-photon calcium imaging (Fig. 4, 6). In contrast to modeling predicting poor entrainment of cell bodies proximal to the electrode [22], proximal and distal neurons were activated suggesting selective activation of distal neurons was not achieved (Fig. 4a, 6). In fact, the populations neurons activated by symmetric cathodic and asymmetric anodic stimulation were nearly identical (Fig. 8). It is likely fibers of proximal neurons are therefore being activated by the stimulation, possibly resulting in antidromic somatic activation. 10 Hz stimulation was chosen to limit the effect of inactivation distal neurons observed during higher frequency stimulation [24, 60], however, it still remains to be determined if the entrainment efficiency at higher stimulation frequencies would be modulated by stimulation waveform.

Additionally, similar to cathodic stimulation, the modulation of activation area could be a result of a difference in activation threshold. If increasing anodic asymmetry does approximate monophasic cathodic stimulation (See 4.1.2), activation at the same current density could result in greater distal activation [75], due to the reduced activation threshold [21]. Although anodic stimulation has been discussed as being more effective at activating cell bodies [27], Voigt *et al.* demonstrated anodic stimulation is much less effective at activating neurons in the auditory cortex [68], which could, therefore, reduce the activation distance. While symmetric anodic stimulation did result in significant reductions in the magnitude of calcium activity (Fig. 5, 7), reductions in the spread of activation were not as pronounced (Fig. 4, 6). This may suggest that waveform shape has a stronger influence on the spatial spread of activation than stimulation amplitude within layer II/III of somatosensory cortex within safety limits. However, it is important to note that increasing the charge per phase beyond 2.5 nC/phase could enter the range of current where safety may be a concern [51]. These observations, therefore, demonstrate how waveform asymmetry can be used to modulate the spatial selectivity of neuronal activation near the maximal limit for safe charge per phase and current density.

### 4.2.3 Spatial selectivity of stimulation asymmetry may depend on cortical location

While the present study demonstrates significant interactions of polarity and asymmetry on the spatial localization of neuronal activation in layer II/III of somatosensory cortex, these effects may differ depending on depth, or cortical location. In fact, asymmetric cathodic leading stimulation in layer IV of the rat barrel cortex did not increase spatial localization of neuronal activation at high amplitudes when measured with voltage sensitive dye imaging [78]. These contrasting results can likely be explained by the difference in the neuronal projection anatomy. Layer IV axonal projections to the superficial layers of cortex is largely restricted to a single cortical column, and layer II/III has greater horizontal projections that can span more than one column [78, 80]. The animal model in this study, which has dense GCaMP6s expression in layer II/III [39], was therefore advantageous to evaluate the horizontal spread of neuronal activation given the intercolumnar expression. However, this difference in fiber orientation could also impact the activation threshold.

Voigt *et al.* demonstrated that the strength of an electrical stimulation evoked response was depth dependent [81], anodic stimulation of infragranular was ineffective at eliciting a cortical response compared to cathodic stimulation [68]. They explained these results citing the assumption that anodic stimulation may be more effective at activating cell bodies, resulting in less activation of proximal neurons whose fibers pass by the electrode. If this is the case, then asymmetric cathodic stimulation would also be less effective at eliciting a cortical response in these deeper layers, at least near the threshold current, which agrees with the reduced magnitude change in voltage sensitive dye reported by Bari *et al [78].* However, it should be noted that in one study in the motor cortex, anodic leading stimulation had lower electrophysiological thresholds than cathodic leading stimulation in deeper cortical layers [82], but in another anodic current had higher thresholds for eliciting movement in deeper layers [83]. Together, given the diversity of cell types and their fiber arborizations among different brain regions and cortical layers, these results highlight the complexity of the mechanisms behind ICMS and emphasize the importance of characterizing the responses of neuronal subtypes.

### 4.3 Stimulation polarity and asymmetry modulate the temporal network dynamics of neuronal activation

Careful control of the temporal dynamics of neuronal activation can be critical important for the application of therapeutic ICMS. Modeling suggests both excitatory and inhibitory input greatly influence the ability for neurons to entrain ICMS [32]. Because this baseline excitability can be impacted by ongoing neuronal activity, it is possible that the activation threshold is also impacted. In fact, Voigt *et al*. demonstrated that ICMS during different phases of neurons activity following an auditory stimulus differentially modulated the network response [81]. Since activation threshold can impact the spatial spread of activation, it is, therefore, likely that the excitatory and inhibitory balance is critical for the spatial spread of activation; however, the impact during long stimulation trials is not entirely clear.

Mesoscale imaging demonstrated that all stimulation waveforms resulted in a substantial reduction in activation magnitude and area (Fig. 2, 3), potentially suggesting poor entrainment of distal neurons to direct or indirect (trans-synaptic) activation. Mechanisms governing poor entrainment of distal neuronal somas may involve ion buildup [31, 84-86], fatigue [87], virtual anode formation [24, 27], or even geometrical branching of axons [30, 88]; however, lateral inhibition may play the largest role [75]. Electrical stimulation in cortical slices demonstrated lateral inhibition is critical for containing the horizontal spread of neuronal activation in the cortex [89]. Although this reduction in somatic entrainment distal to the electrode is more prevalent at higher frequency [24], the presence of this effect at low frequency in all waveforms suggests an important influence of other neuronal cell types.

To further support this claim, mesoscale imaging revealed a prolonged substantial reduction in fluorescence intensity following stimulation, which was particularly evident for the waveforms resulting in the largest spatial spread of activation (Fig. S3). A greater spread of activation would be likely to recruit greater inhibitory input, either directly or indirectly, therefore resulting in greater reduction of neural activity following stimulation. However, if it is inhibition, it is unclear why this effect was not observed during two-photon imaging. While the size of the imaging plane may influence the ability to observe this effect (407 μm vs. 3 mm), mesoscale imaging averages the neuronal activity of several layers, and a reduction in direct or indirect layer I neuronal activity without a reduction in layer II/III activity could still result in a decreased activation area. Furthermore, since astrocytes are key regulators of neurotransmitter clearance, and astrocytic voltage changes can influence the efficiency of this regulation [90] they could also influence the entrainment of somas. Investigation into these effects may be even more important in the context of temporal patterned stimulation [91], in which different neuronal subtypes can differentially entrain during the pulse bursts likely modulating the excitatory inhibitory balance during the inter-pulse intervals. Ultimately, multimodal imaging allowed further insight into the spatial dynamics of neural activation and the role waveform plays in shaping recruitment of different neurons.

### 4.4.1 Asymmetric cathodic waveforms as an effective tool in neural prosthetics

Cathodic leading asymmetric stimulation was hypothesized to selectively activate cell bodies orthodromically [21], thus resulting in spatially localized activation of neurons around the electrode. This could be particularly useful to achieve highly localized sensory percepts in cortical activation, or in spinal cord stimulation where fibers and cell bodies of different circuits are in close proximity. This study demonstrates that the benefit of spatial localization with asymmetric cathodic pulses (Fig. 2, 4, 6), comes at a cost of reduced activation magnitude at the same current density (Fig. 3, 5, 7). Consistent with this, multiple studies showed that the behavioral detection threshold for cathodic first asymmetric stimulation is increased compared to symmetric cathodic stimulation [4, 42]. Therefore, increased current will likely be required to achieve the same perceptual intensity even if it is localized. Although, consideration of the layer of stimulation, and the level of cortical activity could impact the spatial spread and magnitude of activation [79, 81]. Importantly, however, the effect of different waveforms on perceptual quality may be an important outcome to consider.

The temporal rate code of neural activation is an important dimension in sensory perception [92]; therefore, the modulation of the temporal activation of diverse neuronal populations with different waveforms could increase the dynamic range of artificial perception. Increasing cathodic asymmetry increased the activation time to peak of neuronal activation, the rate of rise of calcium activity (Fig. S1, S2), and the activation time of the activated neurons (Fig. 7). There was also a greater percentage of “delayed response” cells in the population of activated neurons (Fig. S3). This may suggest that these waveforms may be able to encode a perceptual stimulus that can increase over time. Alternatively, these waveforms could be beneficial for therapeutic applications requiring selective activation during long stimulation trains, such as deep brain stimulation.

### 4.4.2 Asymmetric anodic waveforms as an effective tool in neural prosthetics

Asymmetric anodic stimulation was hypothesized to be beneficial in spinal cord stimulation to selectively activate passing fibers without entrainment of the local cell bodies [22]. The observation that the neural populations activated by asymmetric anodic waveforms was nearly identical to those activated by symmetric cathodic (Fig. 8), may explain why behavioral detection thresholds were comparable [18]. Although proximal cell bodies are activated in ICMS of layer II/III, it is not clear how the entrainment of these neural populations may change at higher frequencies. In fact, the majority of neurons preferred the most asymmetric anodic stimulation investigated (A_7_; Fig. 8). Because fiber geometry, frequency, and the excitatory and inhibitory balance greatly influence the level of entrainment [32], it will be interesting to investigate how waveform modulates these effects to better inform clinical therapies employing electrical stimulation.

### 4.5 Limitations

This study used *in vivo* calcium imaging in male mice to address the hypothesis that stimulation waveform can modulate the spatial distribution and entrainment of neurons around an intracortical electrode. The guiding principle behind the use of asymmetric waveforms is that the long duration leading phase can modulate the activation probability of different neuronal elements and lead to either selective orthodromic (cathodic first) or antidromic (anodic first) activation. Due to the kinetics and spatial localization of the calcium sensor used in this study, we are unable to confirm the location of stimulation initiation. Although we interpret greater spread of activation to be an increased probability of antidromic activation, we cannot separate direct synaptic activation from indirect activation through synaptic transmission. Furthermore, although we were primarily interested in characterizing how waveform affected activation given the same current density, activation threshold is affected by stimulation polarity [68] and waveform shape [21]. Therefore, the spatial modulation could be a result of sub or suprathreshold activation, but because the current density employed in this study is close to the safety limit of maximal current [51] and the range of behavioral detection thresholds for human somatosensory stimulation [4], these results are likely informative for clinical investigation.

Additionally, because the calcium sensor used in this study (GCaMP6s) increases fluorescence with the number of action potential in short succession [39], we defined entrainment to refer to the ability of neurons or neuropil to result in activation to greater, or lesser percentage of stimulation pulses as measured by a larger increase in fluorescence. Importantly, it is not clear whether the increased fluorescence intensity is a result of greater 1:1 entrainment, or due to other activation properties such as bursting. Newer developments in genetically encoded voltage (GEVI) and calcium indicators (GECI) with faster kinetics may be able to more comprehensively address these questions [93–95]. However, it should be noted that current GEVIs do not provide comparable spatial resolution and suffer from lower signal to noise ratio [96]. Therefore, the relative magnitude of calcium activity likely still provides valuable information about the level of stimulation-induced neuronal activity.

While highly valuable for spatial resolution, due to the limitations of two-photon imaging, we could only observe direct neuronal activation within the imaging plane. Newer developments in light sheet [97], or even swept confocally-aligned planar excitation (SCAPE) imaging [98] could be suited to observe not only the horizontal activation, but also the activation between cortical layers at high speeds. These tools could be particularly useful in determining how waveform may affect neuronal activation in different cortical layers especially considering polarity can differentially affect neuronal activation depending on the depth [68]. Still, these results are informative for clinical ICMS because the depth of electrode may vary during implantation, and it is likely that some of the sites of a Utah array remain in layer II/III [99].

Lastly, this study was conducted in an acute preparation necessitating the use of anesthesia. The use of ketamine, a known NMDA antagonist and GABA_A_ potentiator, likely modulated the baseline probability of neuronal activation and entrainment [100–102]. Future studies should be conducted in awake preparations, taking particular care to consider the large influence on neuronal damage, microglial activation, and astroglia scarring, which are suggested to modulate both recording and stimulation [67, 103, 104]. Additionally, there is a need to directly evaluate how stimulation induced activity changes over time in a chronic setting given the dynamic tissue responses to implantable electrodes and its effect on the excitatory inhibitory balance.

### 4.6 Conclusion

Using calcium imaging, we demonstrated that stimulation waveform and polarity largely interact in the spatial recruitment of neurons. As asymmetry index increases in cathodic first pulses, spatial localization increases, and level of neuronal activation decreases. Conversely, in anodic first pulses, the activity level and spatial distribution begins to better resemble conventional symmetric cathodic leading stimulation. Even at 10 Hz, there is a spatial fall off of neuronal activity to stimulation over a 30s train. Additionally, there is a prolonged period of inactivity following stimulation that scales with the level of activation during stimulation. Importantly, this demonstrates the potential for stimulation waveform in modulating neuronal activity for different therapeutic applications, or even elicitation of complex sensory percepts. Furthermore, with the widespread use of electrical stimulation in circuit analysis, we provide evidence of stimulation parameters that can increase spatial selectivity and limit the activation of passing fibers from different circuits. Thus, stimulation polarity and asymmetry can, therefore, be used to investigate the role of neuronal entrainment and activity within a target circuit.

## Author Contributions

*Conceptualization*, All authors; *Methodology*, All authors; *Investigation,* K.C.S., *Formal Analysis*, K.C.S. and J.R.E. *Writing*—*Original Draft*, T.D.Y.K., and K.C.S.; *Writing*—*Review & Editing*, All authors; *Visualization*, K.C.S and T.D.Y.K.; *Supervision*, T.D.Y.K.; *Funding Acquisition*, T.D.Y.K.

## Acknowledgments

This work was supported by NIH R01NS094396, R01NS105691, and R21NS108098, NSF CAREER 1943906, and the ARCS Foundation, Inc., Pittsburgh Chapter, Gookin Family Foundation Award.

## Data Accessibility

The data that support the findings of this study are available from the corresponding author upon reasonable request.

---

**Figure S1:**
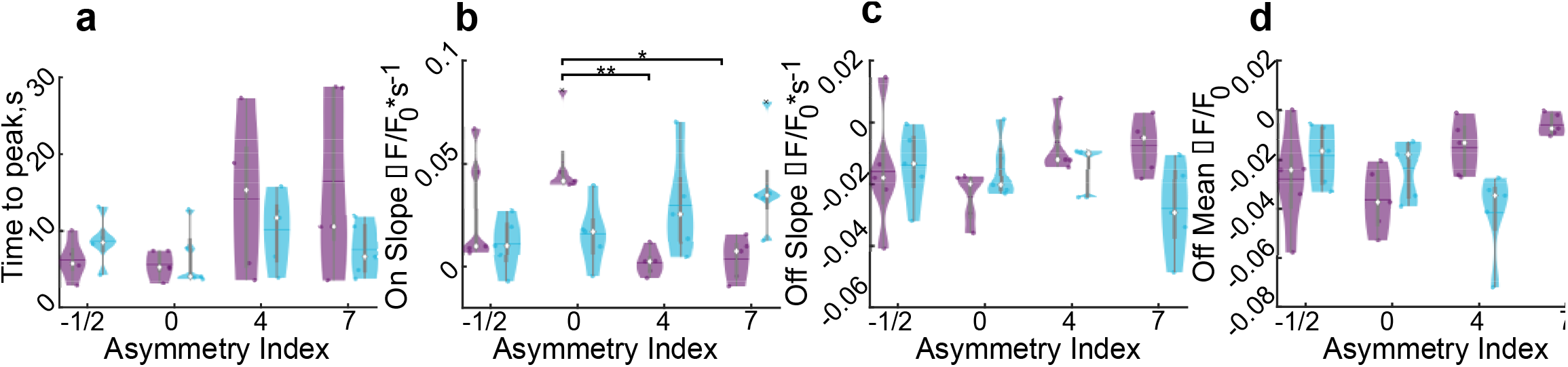
Mesoscale activation timing. (a) Time to peak activation over 980 μm during stimulation. (b-c) The activation (i; F_3,12_=9.16, n=5, p=0.002) and inactivation (j) slopes of ΔF/F_0_ over 980μm. (d) The mean ΔF/F_0_ of the 10s post stimulation averaged over 400 μm shows substantial reduction from baseline in all conditions (interaction F_3,12_=8.38, n=5, p=0.0028).Statistics were assessed with 2-way rmANOVA with post-hoc Welch’s T-test and Bonferroni-Holm correction. * indicates p<0.05, ** indicates p<0.01.

**Figure S2:**
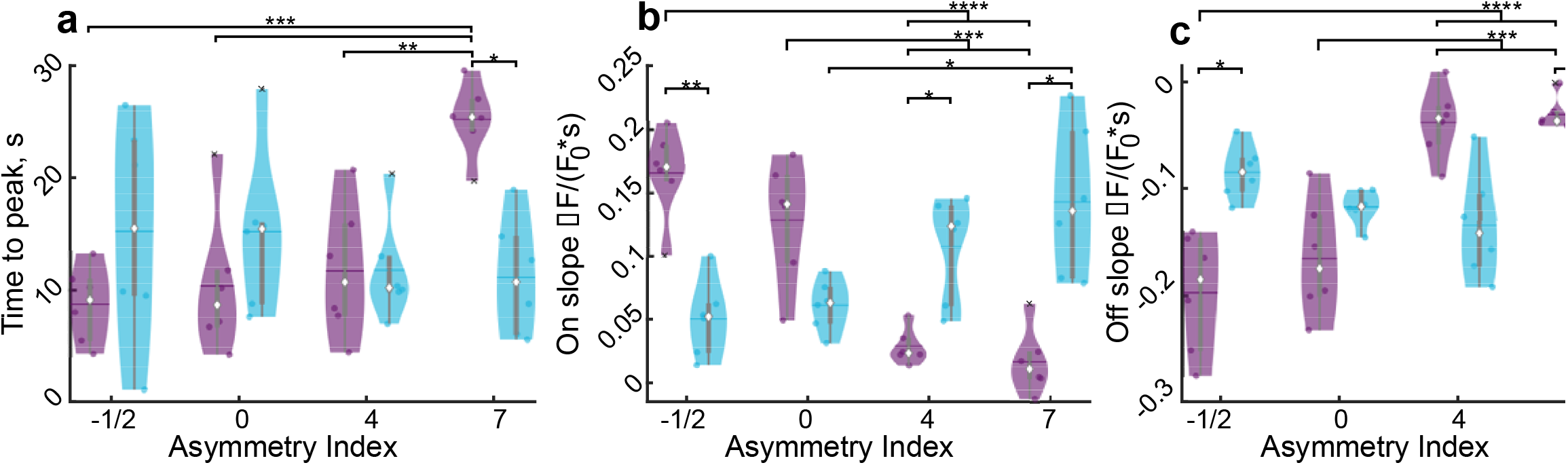
Neuropil activation timing. (a) Time to peak activation is increased in C_7_ (h, F_3,15_=5.06, n=6, p=0.0128), the rate of change of fluorescence during the first 2 seconds of stimulation (I; F_3,15_=24.47, n=6, p<1e−5) and the last two seconds of stimulation (j; F_0.70,3.52_=21.15, n=6, p=0.0049) are modulated by stimulation waveform. Statistics were first assessed with a 2-way rmANOVA followed by post-hoc Welch’s T-tests with Bonferroni-Holm correction : * indicates p<0.05, ** indicates p<0.01, *** indicates p<0.001.

**Figure S3:**
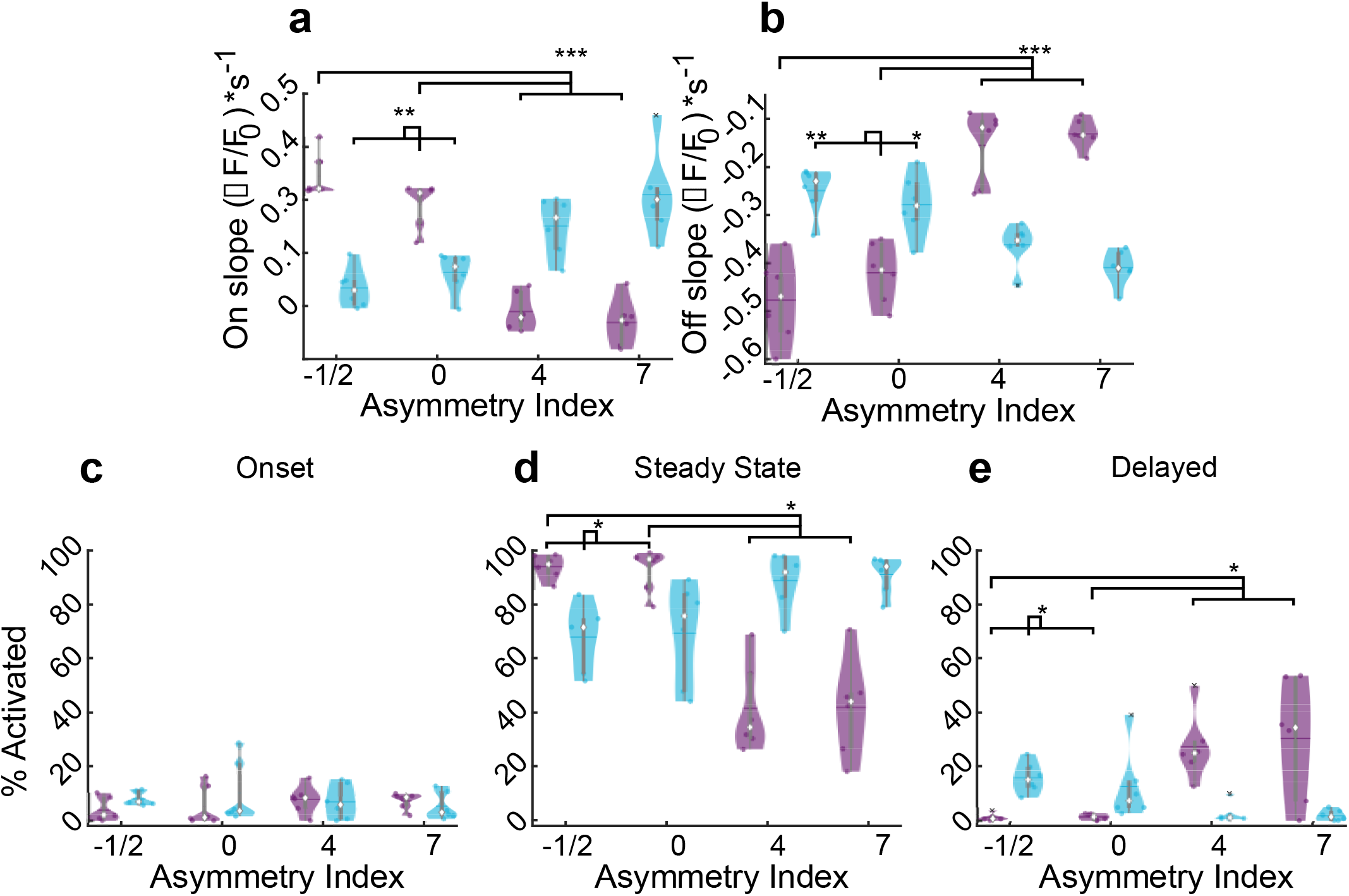
Activation timing of neurons is modulated by stimulation waveform and asymmetry. (a-b) The rate of change of the fluorescence during the first 2 seconds (a; F_3,15_=42.91, n=6, p<1e−6) and last two seconds following stimulation (b; F_0.72,3.61_=44.62,n = 6, p<0.001) are modulated by stimulation waveform. (c-e) Mean percent of cells activated in each animal characterized as onset (c), steady state (d, F_3,15_=56.35, n=6, p<1e−7), or delayed (e; F_3,15_=22.41, n=6, p<1e−5). Statistics were first assessed with a 2-way rmANOVA followed by post-hoc Welch’s T-tests with Bonferroni-Holm correction : * indicates p<0.05, ** indicates p<0.01, *** indicates p<0.001.

